# A global view of meiotic double-strand break end resection

**DOI:** 10.1101/067199

**Authors:** Eleni P. Mimitou, Shintaro Yamada, Scott Keeney

## Abstract

The DNA double-strand breaks that initiate homologous recombination during meiosis are subject to extensive 5′→3′ exonucleolytic processing. This resection is a central and conserved feature of recombination, yet its mechanism is poorly understood. Using a purpose-made deep-sequencing method, we mapped meiotic resection endpoints genome-wide at high spatial resolution in *Saccharomyces cerevisiae*. Generating full-length resection tracts requires Exo1 exonuclease activity and the DNA-damage responsive kinase Tel1, but not the helicase Sgs1. Tel1 is also required for efficient and timely initiation of resection. We find that distributions of resection endpoints at individual genomic loci display pronounced heterogeneity that reflects a tendency for nucleosomes to block Exo1 in vivo, yet modeling experiments indicate that Exo1 digests chromatin with high apparent processivity and at rates approaching those for naked DNA in vitro. This paradox points to nucleosome destabilization or eviction as a determining feature of the meiotic resection landscape.

## Main Text

Meiotic recombination plays a pivotal role in sexual reproduction by promoting proper segregation of homologous chromosomes^1^. Recombination initiates with DNA double-strand breaks (DSBs) formed by the topoisomerase-like transesterase Spo11, which remains covalently attached to DSB 5′ ends (Fig. 1a). Endonucleolytic cleavage dependent on the conserved Mre11– Rad50–Xrs2 (MRX) complex in cooperation with Sae2 generates nicks on the Spo11-bound strands that serve as entry points for the modest 3′→5′ exonuclease activity of Mre11 and the much more robust 5′→3′ exonuclease activity of Exo1 (refs 2,3–5). The net result is the release of Spo11 bound to a short oligonucleotide (oligo) and, more importantly, generation of long 3′ single-stranded DNA (ssDNA) tails. These tails are substrates for the strand-exchange proteins Dmc1 and/or Rad51, which carry out homology search and invasion of a repair template in the homologous chromosome^1^. A similar nick-plus-exonuclease mechanism operates in vegetative cells^6^, except that Exo1 is partially redundant with the concerted action of the Sgs1–Top3–Rmi1 complex along with the Dna2 nuclease for extensive 5′→3′ resection^7,8^. In meiosis, available data clearly implicate Exo1 but suggest that Sgs1 is dispensable^4,9^.

**Figure 1:**
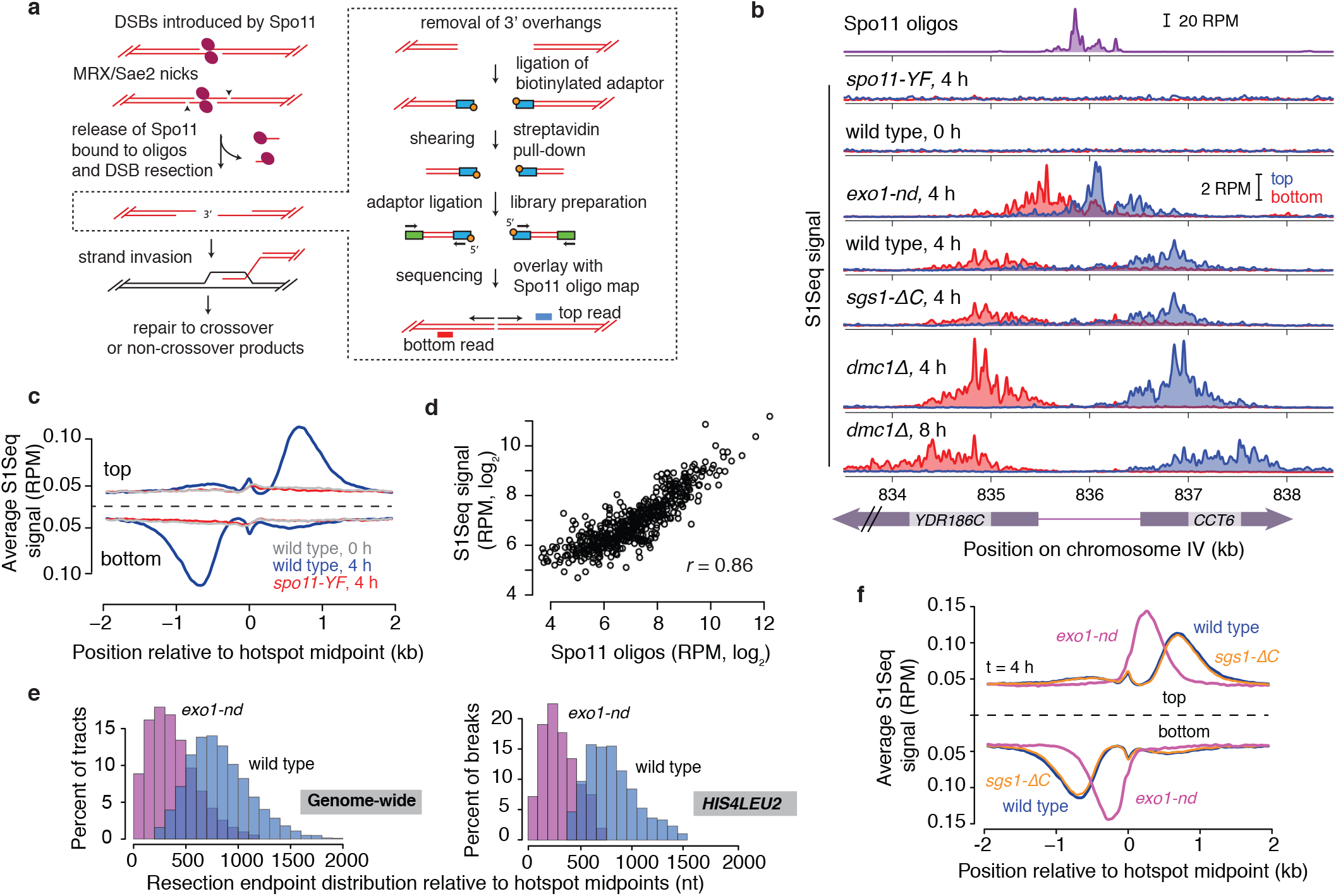
Genome-wide mapping of meiotic DSB resection. **a**, Schematics of early steps in recombination and of the S1Seq assay (boxed). DNA from meiotic cells was embedded in agarose plugs to protect from shearing, digested with S1 nuclease and T4 DNA polymerase, and ligated to a 5′-biotinylated dsDNA adaptor. After extraction from plugs and sonication, biotinylated fragments were affinity purified, ligated to a second adaptor, amplified by low-cycle PCR, deep sequenced, and mapped to the yeast genome. **b**, Snapshot of DSBs (Spo11-oligo sequencing^13^) and strand-specific S1Seq reads surrounding the *CCT6* hotspot in different genetic backgrounds (51-bp smoothing window). *spo11-YF* is the catalytically inactive *spo11-Yl35F* mutant; *sgsl-ΔC* expresses a C-terminal truncation lacking the helicase domain. **c**, Genome averages of strand-specific S1Seq signal (101-bp smoothing) centered on midpoints of Spo11-oligo hotspots (n=3908). **d**, S1Seq signal correlates well with Spo11-oligo counts. S1Seq signal (wild type, 4 h) was summed in the regions from +200 to +1600 bp (top strand) and –200 to –1600 bp (bottom strand) relative to hotspot midpoints. Only “loners” were considered, i.e., no other hotspot within 3 kb. **e**, Comparison of the genome-wide distribution of resection endpoints from S1Seq with resection lengths measured by Southern blotting at the artificial *HIS4LEU2* hotspot^4^. **f**, Genome averages as in panel c.

Although meiotic resection was first demonstrated experimentally more than 25 years ago and has long been known to be a fundamental step in recombination^10–12^, it remains poorly understood. In fact, high resolution, quantitative data are only available for one side of a single artificial DSB hotspot^4^ and whether this information can be extrapolated to natural DSB hotspots genome-wide is unknown. To overcome this limitation, we developed methods to map resection endpoints genome-wide specifically and with high sensitivity and spatial resolution. These maps allowed us to answer longstanding questions about the genetic control of resection; the handoff from MRX–Sae2 to Exo1; locus-to-locus variation in fine-scale resection patterns; the kinetics of resection; and how resection machinery interacts with barriers imposed by local chromatin structure.

## Mapping DSB resection endpoints

We digested the 3′ tails of resected DSBs with ssDNA-specific nucleases to generate sequencing libraries of ssDNA–dsDNA junctions that were then compared to previously derived DSB maps from Spo11-oligo sequencing^13^ (Fig. 1a,b and Extended Data Table 1). For each strain or time point, data were pooled from two highly reproducible biological replicates (Pearson’s *r* = 0.95–0.99) (Extended Data Fig. 1a,b).

Reads from resection endpoints should be meiosis-specific and Spo11-dependent and should flank DSB hotspots with defined polarity (Extended Data Fig. 1c). Indeed, S1 sequencing (S1Seq) reads were enriched with the expected polarity around hotspots, and this enrichment was absent from premeiotic (0 h) samples and from meiotic samples from the catalytically inactive *spo11-Y135F* mutant (Fig. 1b,c and Extended Data Fig. 1d,e,f). S1Seq signal around hotspots correlated well with Spo11-oligo counts (Fig. 1d), and reads spread further from hotspots in a time-dependent manner in *dmc1Δ* (Fig. 1b and Extended Data Fig. 1e,g,h), consistent with gradual hyper-resection^14^. We conclude that S1Seq is a sensitive and quantitative measure of DSB resection endpoints.

Resection endpoints were located from ~200 to ~2000 nt away from hotspot centers, with a mean of 822 nt and a positive skew (Fig. 1c,e). This pattern resembles Southern blot analysis of the *HIS4LEU2* hotspot^4^ (Fig. 1e, range 350–1550, mean 800 nt), but greater sensitivity of S1Seq captured less abundant species at extremes of the distribution. Genome-average profiles were smooth and left–right symmetric (Fig. 1c), but individual hotspots were heterogeneous with strong peaks and valleys that differed between hotspots and between sides of the same hotspot (Fig. 1b and Extended Data Fig. 1e). This heterogeneity reflects previously undescribed effects of chromatin structure on resection termination (see below).

S1Seq profiles affirmed dispensability of Sgs1 (Fig. 1b,f and Extended Data Fig. 1e). In contrast, the nuclease-defective mutation *exo1-D173A* (*exo1-nd*)^15^ caused S1seq reads to cluster closer to DSB hotspots, reducing resection lengths to <1100 nt (mean = 373 nt), comparable to *HIS4LEU2* in *exo1*Δ (mean = 270 nt)^4^ (Fig. 1b,e,f and Extended Data Fig. 1e). We consider it likely that the distribution of resection endpoints in *exo1-nd* is also the distribution of the most distal Mre11-dependent nicks that form in wild type. If so, the difference between Spo11-oligo lengths and *exo1-nd* endpoints indicates that there are multiple Mre11-dependent nicks on the same strand, or that there is a single distant nick plus extensive 3′→5′ digestion averaging 335 nt (Extended Data Fig. 1i).

## Tel1 promotes initiation and extension of resection

Absence of Tel1 (a DSB-responsive kinase orthologous to human ATM) decreased resection length at *HIS4LEU2* for early DSBs^16^. This and other findings led to the proposal that Tel1 controls resection when DSB numbers are still low, whereas higher DSB numbers later allow Mec1 (ATR in humans) to substitute^16^. S1Seq data refined this model and revealed that Tel1 acts at multiple steps in resection.

S1Seq reads fell closer to DSB hotspots in the *tel1Δ* mutant at both early (2 h) and later (4 h) time points, indicating shorter resection, but with peaks in similar positions as in wild type (Fig. 2a,b). Although a subset of tracts at 4 h in *tel1Δ* matched the longest in wild type (Fig. 2b), DSBs remained hypo-resected in the mutant even at this later time (Fig. 2a,b), so Tel1 influences resection length throughout meiosis.

**Figure 2:**
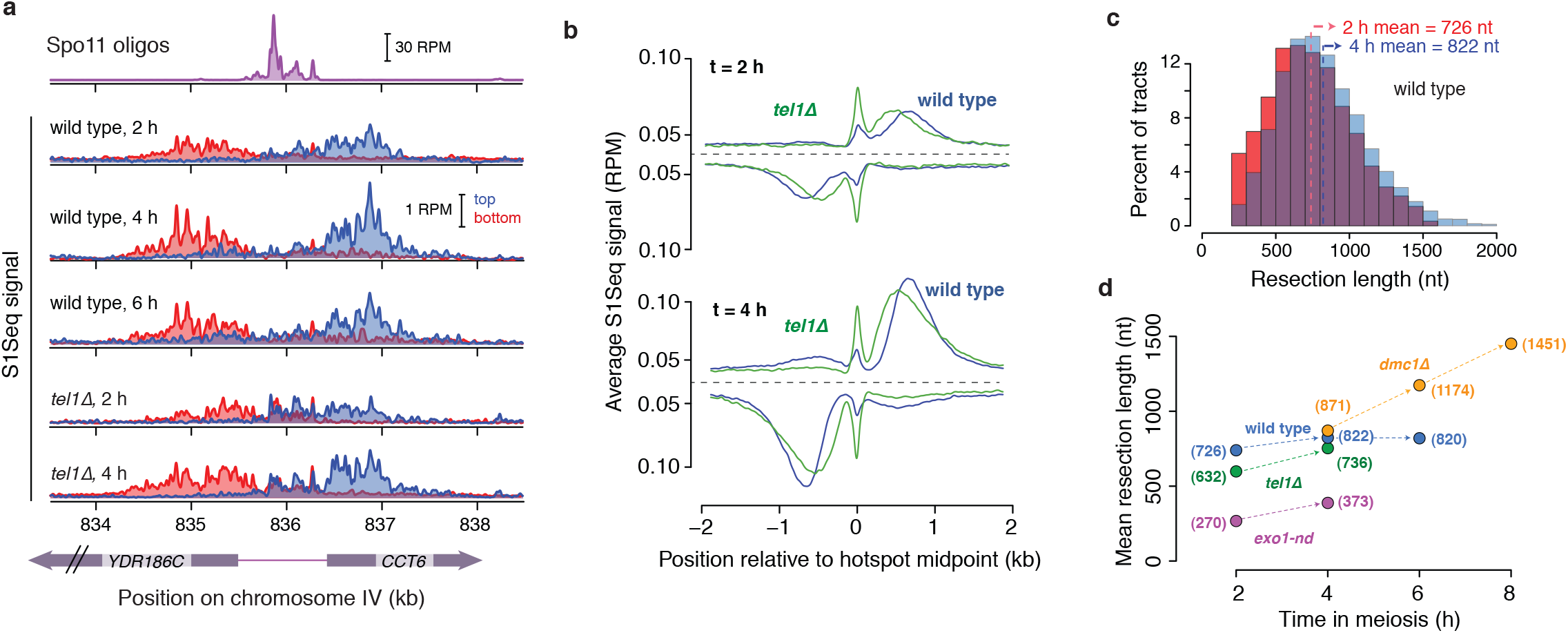
Tel1 and the evolution of resection tract lengths over time in meiosis. **a**, S1Seq (51-bp smoothing) at *CCT6* over meiotic time courses in wild type and *tellΔ*. **b**, Genome averages as in Fig. 1c. **c**, Resection tracts are longer on average later in wild-type meiosis. Data are for loner hotspots. **d**, Mean resection tract lengths as a function of time in meiosis for the indicated genetic backgrounds.

DSBs in wild type have appeared to be maximally resected as soon as they are detectable^14,16^. S1Seq profiles at individual hotspots seemed stable over time in wild type (Fig. 2a), especially compared to *dmc1Δ* (Fig. 1b), but genome-wide averages revealed slightly shorter resection tracts at 2 h than at 4–6 h (Fig. 2b,c,d and Extended Data Fig. 1j). Locus-to-locus variability and the greater sensitivity of S1Seq compared to Southern blotting are nonexclusive possibilities for why this small change was not seen before. Although shorter tracts may reflect partially resected DSBs, we favor the interpretation that later-forming DSBs tend to be resected further, perhaps via increasing Tel1 activity^16^.

Unexpectedly, *tel1Δ* cells had higher S1Seq signal within hotspots, accounting for a greater fraction of reads at 2 h than 4 h in *tel1Δ*, but also present in wild type at lower levels (Fig. 2a,b and Extended Data Fig. 1e). We hypothesized that this signal might reflect presence of unresected DSBs. Within-hotspot S1Seq signal displayed peaks overlapping strong Spo11-oligo clusters (Fig. 2a) and it correlated with hotspot strength in both *tel1Δ* and wild type (Extended Data Fig. 2a,b). Fine-scale patterns matched expectation for preferred Spo11 cleavage 3′ of C residues and for the 2-nt 5′ overhang of Spo11 primary cleavage products (Extended Data Fig. 2c,d). We conclude that unresected DSBs are present in wild type at levels too low to detect readily by Southern blotting; higher levels in *tel1Δ* indicate that Tel1 is required for normal resection initiation, possibly via Sae2 phosphorylation^17^. Consistent with this interpretation, a subset of the DSBs in *tel1Δ* mutants detected by Southern blotting are unresected and have Spo11 still covalently attached (M. Neale, personal communication).

## Recombination intermediates

Along with resection signal, there was weaker S1Seq signal with the “wrong” polarity, e.g., top-strand reads mapping to the left of hotspots (Fig. 1b, and Extended Data Fig. 1d,e). This signal was not resection from neighboring hotspots, it correlated with hotspot strength, it was meiosis-specific and Spo11-dependent, and it was essentially absent in *dmc1Δ* (Fig. 1b,c, and 3a; Extended Data Fig. 1e,f,g,h, and 3a). We therefore conclude that this signal derives from S1-sensitive recombination intermediates (RIs), probably displacement (D) loops from strand exchange (Extended Data Fig. 3b,c). Throughout this study, resection profiles were corrected by subtracting an estimate for the small amount of RI signal with the “correct” polarity that presumably masquerades as resection signal (Extended Data Fig. 3b,c; see Methods).

**Figure 3:**
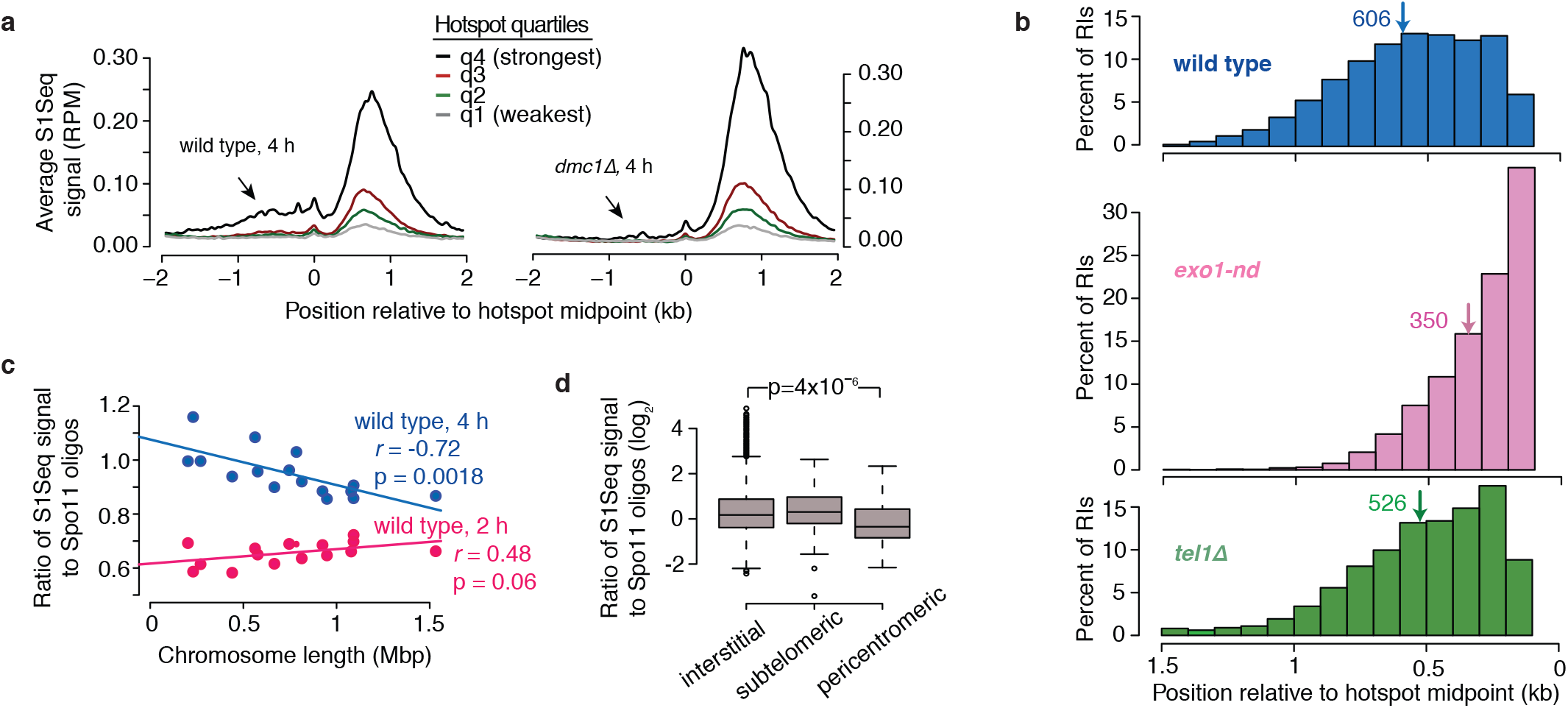
Recombination intermediates detected by S1Seq. **a**, S1Seq RI signal correlates with hotspot heat and is Dmc1-dependent. S1Seq profiles are shown (101 bp smoothing, top and bottom reads co-oriented and averaged) for loner hotspots divided into four groups based on Spo11-oligo counts. Arrows point to region containing RI signal. **b**, Distribution of RI signal relative to hotspot midpoints. Arrows indicate means (bp). **c**, Evidence for longer DSB lifespan on smaller chromosomes. The yield of S1Seq signal (summed as in Fig. 1d) per DSB (Spo11-oligo count) correlates negatively with chromosome size at 4 h but not 2 h in meiosis. **d**, Yield of S1Seq signal per DSB is lower for pericentromeric hotspots (n=82) compared to subtelomeric (n=60) or all other hotspots (interstitial, n=3744). Boxplots are as defined in Extended Data Fig. 2c except that individual points indicate outliers.

We explored spatial and temporal patterns and genetic control of these RIs. At 4 h in wild type, RI reads had a positively skewed distribution with an estimated mean of 606 bp from hotspot centers, i.e., ~25% shorter than the mean resection length (Fig. 3b). The *sgs1* mutant was indistinguishable from wild type (Extended Data Fig. 3d), but RI distances were shorter in *tel1Δ* consonant with the modestly shorter resection tracts (Fig. 3b), suggesting that resection length influences RI position. In *exo1-nd* the RI signal was even closer to hotspot centers (Fig. 1f, 3b, and Extended Data Fig. 3e). The similarity of RI distances (mean of 350 bp) to resection lengths in *exo1-nd* suggests that much of the resection tract is “used up” making these RIs, whereas RIs in wild type usually form within a DSB-proximal subregion of resection tracts.

We reasoned that quantitative differences between Spo11 oligos and S1Seq can reveal regional differences for lifespans of steps in recombination, because turnover of Spo11-oligo complexes is tied to meiotic prophase exit^18^ whereas lifespans of resection and RI signals from S1Seq reflect more directly the progress of the recombination reaction. Smaller chromosomes tend to incur more DSBs per kb because of negative feedback regulation of DSBs tied to engagement of homologous chromosomes^13,18^. It was proposed that smaller chromosomes take longer on average to engage homologous partners because multiple DSBs per chromosome are usually needed for successful engagement^18,19^. This hypothesis predicts that resected DSBs should tend to persist longer on smaller chromosomes. Consistent with this prediction, the ratio of S1Seq resection signal to Spo11-oligo counts at 4 h correlated negatively with chromosome size, i.e., the yield of resected DSBs (S1Seq) per DSB formed (Spo11 oligos) was higher on smaller chromosomes (Fig. 3c). This anticorrelation was not seen at 2 h (Fig. 3c), as expected because early DSBs have not had time to progress further in recombination regardless of chromosome size. The ratio of RI signal to Spo11-oligo counts was uncorrelated with chromosome size (Extended Data Fig. 3f), suggesting that RI lifespan is a function of local features rather than large-scale chromosome pairing kinetics.

The ratio of S1Seq resection reads to Spo11 oligos was lower for pericentromeric hotspots than for other sub-chromosomal domains (Fig. 3d). This apparently shorter lifespan could reflect delayed DSB formation and/or more rapid DSB repair. Consistent with either possibility, kinetochore components suppress DSB formation and promote sister chromatid recombination in pericentric regions^20^.

## Modeling Exo1 mechanism and speed

We tested a model in which Exo1 enters DNA at an Mre11-generated nick and digests DNA until it dissociates from its substrate, which predicts that Exo1 run lengths should follow a geometric distribution defined by the average probability of dissociation at each nucleotide step. We therefore determined the geometric distribution that provided the best fit to wild-type resection lengths when combined with the distribution of presumptive Exo1 entry points (i.e., the *exo1-nd* endpoint distribution) (Fig. 4a). The geometric model fit the data poorly and could be ruled out, but an excellent fit arose when the Exo1 geometric distribution was shifted (Fig. 4a). This simple alternative can be conceptualized in terms of a high probability of Exo1 resecting for a minimum distance (best-fit estimate of 220 nt), after which resection termination follows a geometrically distributed process. This model, plus findings below, provides a framework for understanding the resection mechanism (Conclusions).

**Figure 4:**
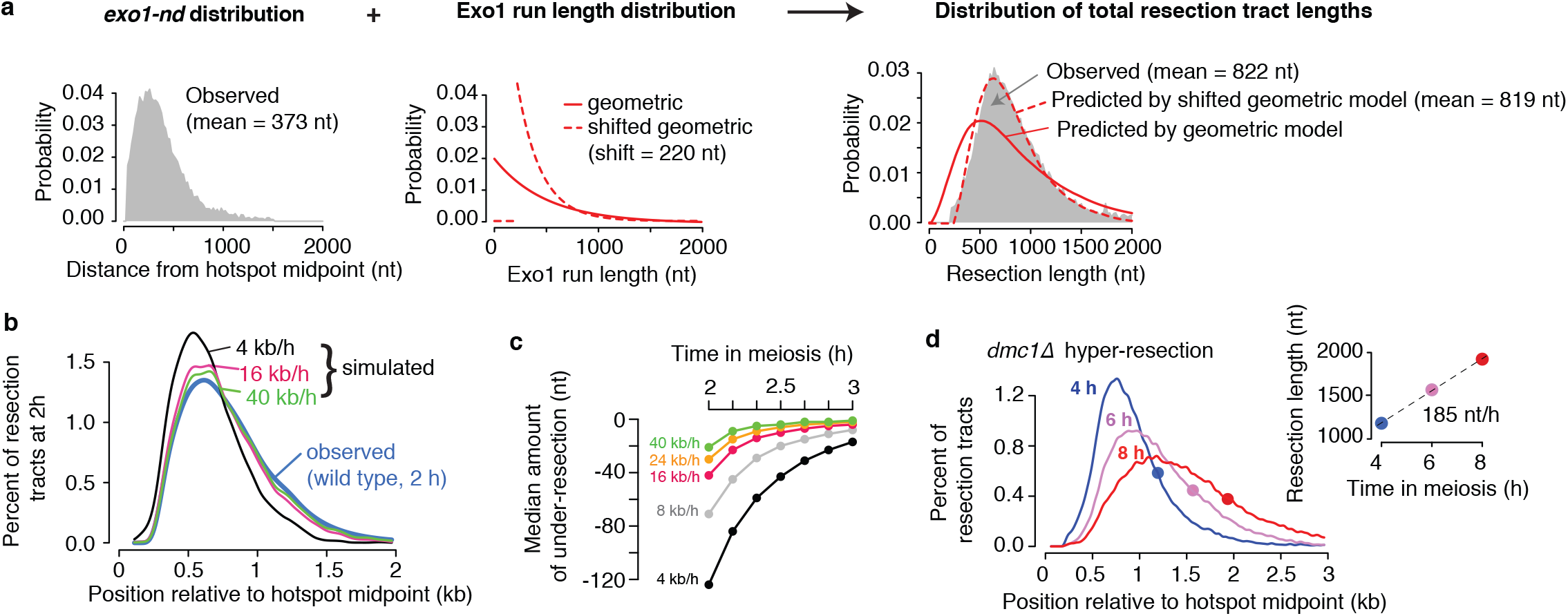
Mathematical modeling of resection. **a**, Modeling Exo1 run lengths. The observed distribution of resection tract lengths (right panel) was modeled by combining theoretical distributions of Exo1 run lengths (middle panel) with the observed distribution of resection endpoints in *exo1-nd* (i.e., a proxy for Exo1 entry points; left panel). Solid red lines: geometric model for Exo1, equivalent to a one-shot processivity model with uniform probability of termination at each exonucleolytic step. Dashed red lines: shifted geometric model for Exo1 run lengths, equivalent to assuming that the probability of Exo1 ceasing resection is essentially zero for some minimum distance. **b,c**, Modeling Exo1 speed. Monte Carlo simulations were used to generate populations of resected DSBs during meiotic time courses, using empirical distributions for the timing of DSB formation^33^ and Exo1 entry points (i.e., *exo1-nd* S1Seq data), plus the shifted geometric model for Exo1 run lengths. Panel b compares simulated and observed resection endpoint profiles at 2 h in meiosis (10-bp binned data, 11-bin smoothing). Note that observed data are fit poorly by an Exo1 rate of 4 kb/h (the vegetative long-range resection rate) because partially resected DSBs are predicted to be a high fraction of total DSBs at this time point. In contrast, rates ≥16 kb/h fit observed data relatively well, indicating that these higher rates are plausible for Exo1 in vivo. Panel c shows the predicted median resection deficit for the simulated resection tracts as a function of time in meiosis for the indicated Exo1 rates. See also Extended Data Fig. 4 and Supplemental Movies 1 and 2. d, Slow hyper-resection in *dmclΔ*. The hyper-resection rate in *dmclΔ* was measured by tracking the leading edge (filled circles; 80th percentile) of resection endpoints (10-bp binned data, 11-bin smoothing; loner hotspots only).

How fast is Exo1? In vegetative cells, unrepairable DSBs are continuously resected for tens of kb at ~4.4 kb/h (refs 8,21,22). Long-range resection by Exo1 only (i.e., in an *sgs1* mutant) is even slower, at ~1 kb/h (ref 8). To test whether the vegetative rate can account for meiotic resection, we performed Monte Carlo simulations to generate populations of resected DNA molecules to compare with S1Seq patterns (Fig. 4b,c, Extended Data Fig. 4 and **Supplemental Movies 1 and 2**. Rates of 4 kb/h or even 8 kb/h were not fast enough: these speeds predicted that DSBs should be resected less far than was observed, particularly at early time points. In contrast, simulations matched observed tracts well when we used rates from 16 to 40 kb/h, the latter being the value from single-molecule studies for Exo1 resecting naked DNA^23^. Since rates ≤8 kb/h appear implausibly slow, we conclude that meiotic resection is much faster than long-range resection in vegetative cells. Because 16 kb/h or higher is plausible, it suggests that Exo1 processes meiotic DSBs in vivo nearly as quickly as it can degrade naked DNA in vitro. Hyperresection in *dmc1Δ* was much slower, with an estimated average speed of 0.19 kb/h (Fig. 4d). Together, these findings clarify the striking differences between the extreme rapidity but limited length of wild-type meiotic resection, the sluggish continuity of hyper-resection in the absence of Dmc1, and the moderate pace but largely unconstrained distance of long-range resection in vegetative cells.

## Chromatin shapes the resection landscape

The strong peaks we observed in S1Seq resection profiles were unexpected. Verifying that this heterogeneity was not a sequencing artifact, S1 treatment converted resected DSB fragments at the *GAT1* hotspot from the usual featureless smears into discrete banding patterns on Southern blots (Fig. 5a and Extended Data Fig. 5a). Prominent bands occupied similar positions in mutants with very different average resection lengths, so this banding pattern is a reproducible feature of the *GAT1* locus itself. S1 generated no discrete banding at *HIS4LEU2* (Extended Data Fig. 5b), consistent with a lack of preferred resection endpoints^4^.

**Figure 5:**
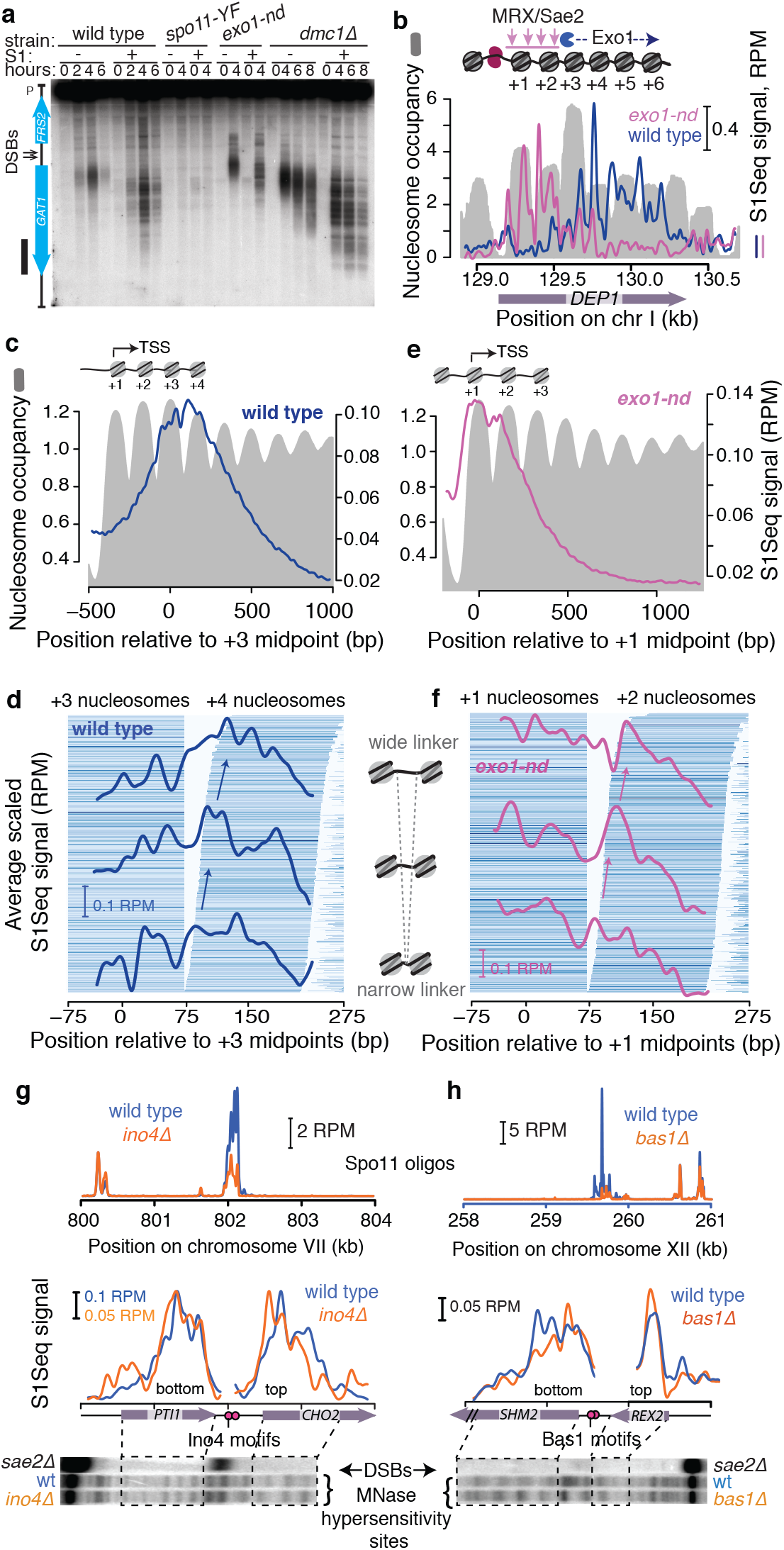
Chromatin structure shapes the resection landscape. **a**, S1 treatment converts the smear of resected DSBs into a discrete banding pattern on Southern blots. **b**, Heterogeneity in resection endpoint distributions within a well-positioned nucleosome array. Top-strand S1Seq reads (51-bp smoothing) are superimposed on a nucleosome occupancy map^13^. Schematic above provides a to-scale illustration of positions of Mre11-dependent clipping and Exo1 digestion. **c**, Average S1Seq resection signal from wild type at 4 h (41-bp smoothed) compared with average nucleosome occupancy, centered on the midpoints of +3 nucleosomes for which there was no other hotspot within 3 kb downstream (n=1815). **d**, Correlation of resection endpoints with nucleosome edges. Pairs of +3 and +4 nucleosomes with robust H3 occupancy (n=962) were rank ordered by increasing linker width. Each horizontal line demarcates a nucleosome position, shaded according to H3 occupancy. Nucleosome pairs were divided into three groups and the averaged S1Seq signal (31-bp smoothed) is superimposed on the nucleosome map. **e,f**, Comparison of *exo1-nd* resection endpoints at 4 h with chromatin structure. Data are presented as for panels c and d, except that averaged profiles are centered on +1 nucleosomes (n=1832 for panel e, n=944 for panel f). **g, h**, Altering chromatin structure alters resection endpoint distributions accordingly. Spo11 oligos and Southern blots of MNase-digested chromatin are from ref. 26.

The stereotyped positions of nucleosomes around natural hotspots allowed us to ask if chromatin structure contributes to this resection heterogeneity. Most yeast promoters have a nucleosome depleted region (NDR), where many DSBs form, flanked by positioned nucleosomes with the transcription start site (TSS) in the first (+1) nucleosome^13,24^ (Fig. 5b). When S1Seq data from wild type were averaged around +3 nucleosomes, the broad peak showed modest scalloping in register with average nucleosome occupancy (Fig. 5c). We speculated that locus-to-locus variation in nucleosome positions blurs the chromatin signature, so we compiled S1Seq averages for three groups of genes divided by linker width between +3 and +4 nucleosomes (Fig. 5d). An S1Seq peak overlapping the left edges of +4 nucleosomes moved progressively to the right with increasing linker width (arrows in Fig. 5d). The widest group also accumulated S1Seq reads within the linkers. Focusing on +2 and +3 nucleosomes revealed similar patterns for the shorter resection tracts in *tel1Δ* (Extended Data Fig. 5c,d), and +1 and +2 nucleosomes yielded similar patterns for Mre11-dependent clipping in *exo1-nd* (Fig. 5e, f). These findings establish a spatial correlation between nucleosome positions and preferred endpoints for both the Mre11 and Exo1 resection steps. For Exo1, the patterns are consistent with digestion proceeding for a short distance into a nucleosome, with increasing likelihood of resection termination as Exo1 approaches the nucleosome dyad possibly reflecting the greater stability of histone-DNA binding near the dyad^25^. RIs displayed a tendency for local peaks at nucleosome linkers (Extended Data Fig. 5e), indicating that RI positions are also influenced by chromatin structure, possibly that of the homologous recombination partner.

To determine if the resection correlation reflects causality, we examined mutants lacking transcription factors Bas1 or Ino4, in which specific hotspots experience altered chromatin structure in adjacent transcription units while retaining promoter-associated DSBs^26^. In *ino4Δ* the nucleosome array in *CHO2* shifts toward the promoter as revealed by micrococcal nuclease (MNase) digestion of chromatin (Fig. 5g). In *bas1Δ* the normally variable nucleosome positioning in *SHM2* (manifesting as a broad, shallow nucleosome ladder in wild type) becomes more regular (Fig. 5h). Concordantly in both cases, S1Seq reads shifted closer to the promoters, whereas neither chromatin structure nor resection in the adjacent *PTI1* or *REX2* genes were much affected (Fig. 5g,h). A control locus where chromatin structure was unchanged in *bas1Δ* showed unaltered resection profiles as well (Extended Data Fig. 5f). Parenthetically, these findings demonstrate that S1Seq heterogeneity cannot be a consequence solely of biases from DNA sequence. More importantly, we can conclude that chromatin structure directly affects Exo1 stopping points, and probably shapes Mre11-dependent clipping as well. To our knowledge, this is the first direct evidence that nucleosome positions determine resection termination positions in vivo.

## Conclusions

Given the typical chromatin structure around yeast DSB hotspots, Mre11-dependent incision often occurs within +1 or +2 nucleosomes and Exo1 then traverses several nucleosomes’ worth of DNA (Fig. 5b). This creates an apparent paradox: the feeble activity of Exo1 on nucleosomal substrates in vitro^27^ and the pronounced tendency for resection to stop near nucleosome boundaries in vivo indicate that nucleosomes are a strong block to Exo1. Yet Exo1 resects for several hundred nt extremely quickly and with high apparent processivity in vivo, as if nucleosomes are initially little or no barrier at all.

A solution to this paradox is for nucleosomes to be destabilized or removed from DNA prior to digestion by Exo1, with a major constraint on resection length simply being how many nucleosomes are removed from Exo1’s path (or are not present to start with) (Extended Data Fig. 5g). Additional factors contributing to resection control may include inhibitors of Exo1, spatially regulated Exo1 activity, and/or iterative Exo1 loading (Extended Data Fig. 5h). Nucleosome eviction at DSBs has been proposed for vegetative yeast and somatic mammalian cells, but experimental support is indirect and conflicting and has generally been unable to distinguish whether observed chromatin changes are prerequisite or consequence of resection^28–30^ (see Supplementary Discussion). Our parameterization of resection speed, apparent processivity, and preferred endpoint positions provides a strong new line of support for this model. Chromatin remodeling enzymes implicated in vegetative resection^31^ are good candidates for nucleosome destabilization in meiotic resection. Interestingly, DSB resection in mouse meiosis extends a similar average distance as in yeast and is inferred to pass through multiple nucleosomes (J. Lange, S.Y., M. Jasin, and S.K., submitted). Conservation of the scale over which DSB-provoked chromatin remodeling occurs could explain this similarity between species.

If nucleosomes are indeed evicted, we can infer that this occurs after DSB formation because Spo11 rarely cuts within nucleosome positions^13^. Furthermore, the strong chromatin signature in the *exo1-nd* mutant implies that the full extent of eviction must occur after positions of Mre11-dependent incision are established. We also note that detection of apparently unresected DSBs in wild type may indicate that Mre11 incision is a rate-limiting step, as in *Schizosaccharomyces pombe^32^*. Tel1 emerges as a prime candidate for promoting Mre11-depended incision given the greater number of unresected DSBs in its absence. Tel1 also regulates net resection distance without changing the positions of preferred stopping points. We speculate that Tel1 controls the efficiency or distance over which nucleosome destabilization occurs (but does not affect translational positions of nucleosomes), which could be via effects on chromatin remodelers, histone modifications, or both.

This study captures the first comprehensive yet finely detailed portrait of the meiotic resection landscape and uncovers components that govern its shape and dynamics. S1Seq gives a selective, sensitive and quantitative measure of DSB resection tracts and should be readily applicable to map resection or DSBs in other settings and organisms.

## Methods Summary

Yeast strains are of the SK1 background (Extended Data Table 2). Synchronized meiotic cultures were prepared according to standard methods. Resection overhang removal took place in agarose plugs with a sequential digestion with S1 nuclease and T4 DNA polymerase, followed by ligation of the first adaptor in plugs. After plug extraction, size selection and shearing, the biotin-containing DNA fragments were affinity purified with streptavidin beads and a second adaptor was ligated. The library was then amplified with low-cycle PCR using primers specific for the adaptors, and submitted for deep sequencing.

**Full Methods** and any associated references are available in **Supplementary Information**.

## Acknowledgments

We thank A. Viale and the MSKCC Integrated Genomics Operation for sequencing and N. Socci and the MSKCC Bioinformatics Core for assistance mapping S1Seq reads. Support for core facilities was provided by the NIH/NCI Cancer Center Support Grant P30 CA008748. We thank M. Neale for sharing unpublished information and N. Hunter and R. Liskay for strains. This work was supported by NIH grants R01 GM058673 and R35 GM118092 (to S.K.). E.P.M. was supported in part by a Fellowship from the Helen Hay Whitney Foundation and S.Y. was supported in part by a Kuro Murase MD-JMSA Scholarship.

## Author contributions

EPM developed S1Seq and performed experiments. SY developed in silico modeling of resection length and speed. EPM and SK designed the study, analyzed data and wrote the paper with contributions from SY.

## Extended Data Figure Legends

**Extended Data Figure 1.**
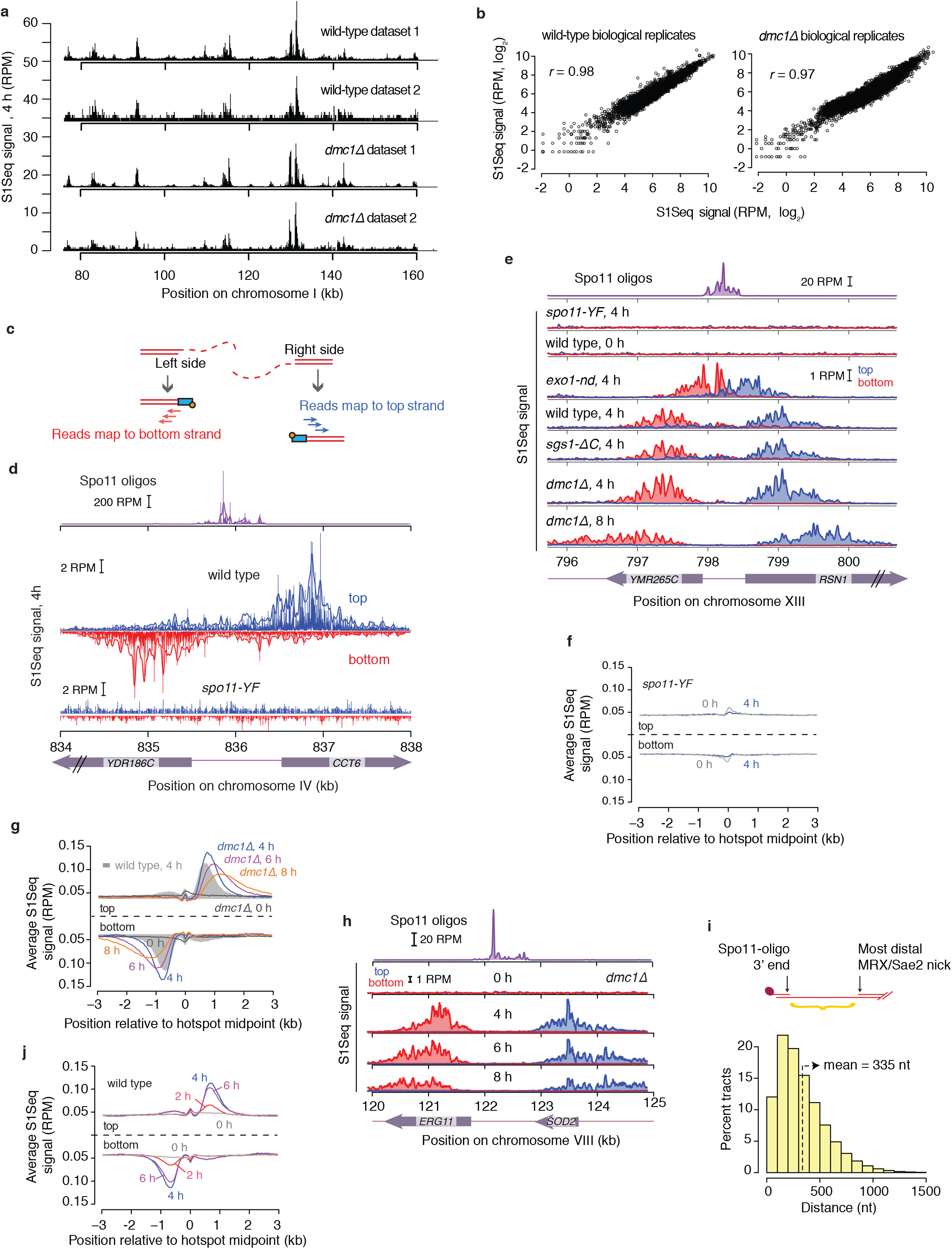
Reproducibility and validation of the S1Seq assay. **a**, Spatial reproducibility of S1Seq maps. S1Seq reads that mapped uniquely to the sacCer2 genome assembly were normalized to reads per million (RPM). A region of chromosome I is shown as an example of the reproducibility between two wild-type and two *dmc1Δ* biological replicates (top and bottom strand reads pooled). Note that apparent differences in background between replicates are caused by differences in sequencing depth (see Extended Data Table 1) and are a trivial visual artifact from overplotting of data points in this relatively large window. **b**, Quantitative reproducibility of S1Seq maps. Representative comparisons are shown for wild-type and *dmc1Δ* biological replicates. Normalized uniquely mapped reads were summed in 1-kb non-overlapping windows. Pearson’s *r* is shown for each. **c**, Schematic of the S1Seq signal polarity. Resection to the right or left should yield reads mapping to the top or bottom strands, respectively. **d**, An example of Spo11-oligo and strand-specific S1Seq reads surrounding the *CCT6* hotspot. Raw data are shown as vertical spikes (RPM value at each bp), with 51-bp smoothing curves superimposed. For clarity, figures in the rest of this paper display only the smoothed data. Note that most of the S1Seq signal displays the polarity expected for resection endpoints, but with an additional minor component of the “wrong” polarity. **e**, A snapshot of Spo11-oligo and S1Seq reads showing another representative hotpot, this time in the *RSN1* promoter, plotted as in Fig. 1b. Note that as at CCT6 (Fig 1b), the *exo1-nd* mutant shortens the spread of the S1Seq signal whereas the *sgs1– ΔC375* mutant lacking the C-terminal helicase domain needed for resection in vegetative cells^8^ is indistinguishable from wild type. **f**, Genome averages for the *spol1-Yl35F* mutant plotted as in Fig. 1c. Note that, while the spread of accumulated S1Seq signal away from hotspot midpoints is absent in *spol1-Yl35F* and therefore DSB-dependent, some residual signal was present at hotspot centers, particularly at 0 h. The source of this signal is not known, but may reflect modest sensitivity to S1 digestion of dsDNA in promoter NDRs tied to typical base and sequence composition unique to these regions^34^. **g**, Genome averages displaying the progressive hyperresection in the *dmc1Δ* mutant. **h**, An illustration of progressive hyper-resection around a representative hotspot in the *dmc1Δ* mutant. **i**, Distribution of distances between the 3′ ends of Spo11 oligos and resection endpoints in *exo1-nd*, which we infer represents the most distal nicks made by MRX plus Sae2. These distances represent either the region over which multiple Mre11-dependent nicks are made, or the lengths traversed by Mre11-dependent 3′→5′ exonuclease activity from a single distal nick back toward Spo11. For this analysis (Fig. 1i) we used the distribution of 3′ ends of mapped Spo11 oligos^13^ and the *exo1-nd* 4 h S1Seq resection reads (10 bp binned) around the midpoints of loner hotspots with width <400 bp corrected for RI signal. **j**, Genome averages over a meiotic time course in wild type.

**Extended Data Figure 2.**
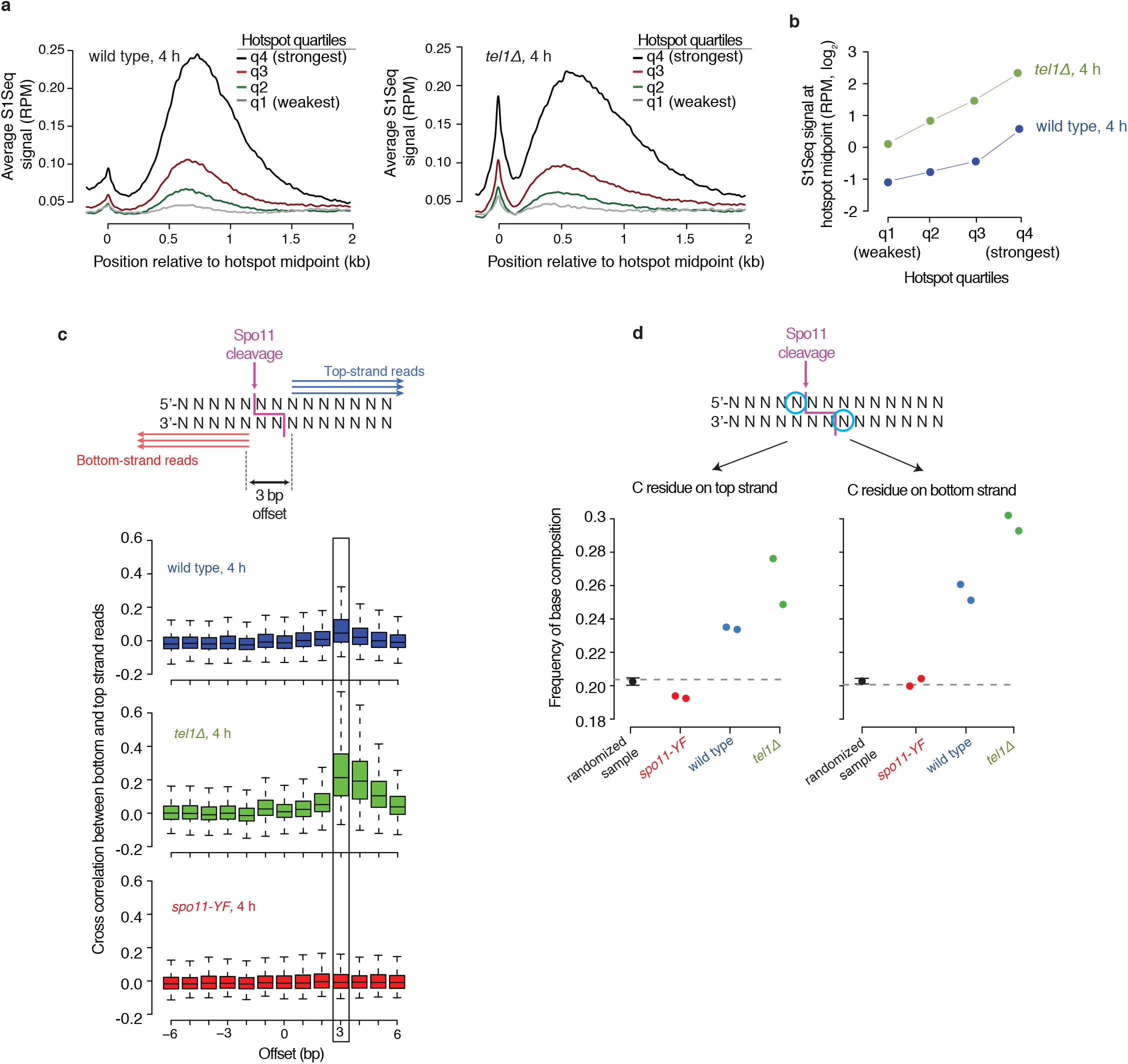
S1Seq detects unresected DSBs in wild type and *tel1 Δ* mutants. **a,b**, Within-hotspot S1Seq signal correlates with hotspot strength. S1Seq profiles are shown in panel a (101-bp smoothing, top and bottom reads co-oriented and averaged) for all hotspots (n=3908) divided into four groups based on Spo11-oligo counts. In panel b, the estimated within-hotspot S1Seq signal correlates with hotspot quartile. We quantified the area under the hotspot-midpoint peak by summing the normalized reads for the region from 0 to 120 bp after subtracting local background (defined as the bottom of the valley between the midpoint peak and resection peak). **c**, Within-hotspot S1Seq signal shows signature consistent with the 2-nt 5′ overhang of unresected DSBs. To determine if within-hotspot S1seq signal might derive from unresected DSBs, we measured the cross correlation between top and bottom strand reads for each of the 711 hottest and narrowest hotspots. Box plots show the distribution of cross-correlation coefficients across this group of hotspots after shifting top and bottom strand reads relative to one another by the indicated offset. Boxes indicate median and interquartile range; whiskers indicate the most extreme data points which are ≤1.5 times the interquartile range from the box; outliers are not shown. Because Spo11 cuts DNA with a 2-nt 5′ overhang^13,35^, overhang removal during library construction should result in top strand reads that are offset by 3 bp from bottom strand reads. Indeed, as predicted, top and bottom strand reads showed a maximal cross-correlation at a 3-bp offset. This cross-correlation was strong in the *tel1Δ* mutant and weaker in wild type, whereas no significant cross-correlation was observed at any offset in *spo11-YF*. **d**, Base composition for S1Seq reads near hotspot midpoints reveals a signature expected for Spo11 primary cleavage products. The DNA sequence of the wild-type, *tel1Δ* and *spo11-YF* 4 h maps was compiled for the regions defined by start and end coordinates for all Spo11 hotspots (n=3908). The plots show the frequency of C residues for the circled positions on top and bottom strands relative to S1Seq reads. Each point shows the result for one biological replicate and the grey dashed line indicates the average C residue frequency in the queried genomic space. As a control, we measured the C frequency for a randomized sample, compiled by randomly selecting 60 positions (10 for each replicate and background) in a 201-bp window around the 5′ ends of the S1Seq reads. The black dots indicate median frequency and the error bars 95% confidence intervals for the randomized control. Spo11 cleavage is favored 3′ of C residues^13,36^. If a subset of within-hotspot S1Seq reads reflect unresected DSBs, this Spo11 cleavage preference predicts concordant nonrandom base composition at the 5′ ends of S1Seq reads. Indeed, we observed base composition enrichment consistent with this prediction in wild-type and more strongly in *tel1Δ*, but not in *spo11-YF*. ***Notes***: We conclude from these analyses that S1Seq is detecting unresected DSBs in wild type and in *tel1Δ* mutants. In panels c and d, the signatures are weaker for wild type than *tel1Δ*, from which we infer that unresected DSBs are relatively frequent in *tel1Δ* but rare in wild type. We consider it likely that in both genotypes these are DSBs that would eventually have been resected, but it is possible that they reflect transient Spo11 cleavage events that are destined instead for resealing (assuming Spo11-mediated DNA cleavage is reversible, as for other topoisomerase family members). In principle, two cuts of the same DNA molecule could release the intervening fragment still bound to Spo11, without intervention by Mre11-dependent incision. Such doubly-cut molecules would be expected to be more frequent in *tel1Δ* mutants because Tel1 prevents an inherent tendency for such multiple cutting to occur^37^. However, even if such molecules are formed, it is unlikely that they would contribute significantly to our sequencing libraries because the size selection step we perform in between ligation of the first and second adaptors would remove them. We note two further implications. First, we could have detected these signatures of unresected DSBs only if S1Seq often (if not always) retains single-nucleotide resolution for mapping ssDNA–dsDNA junctions. We therefore infer that maps of resection endpoints are similarly resolved at this high resolution. Second, our findings indicate that S1Seq can provide a novel and simple method for nucleotide-resolution mapping of the unresected DSBs that accumulate in *sae2Δ* or *rad50S* mutants.

**Extended Data Figure 3.**
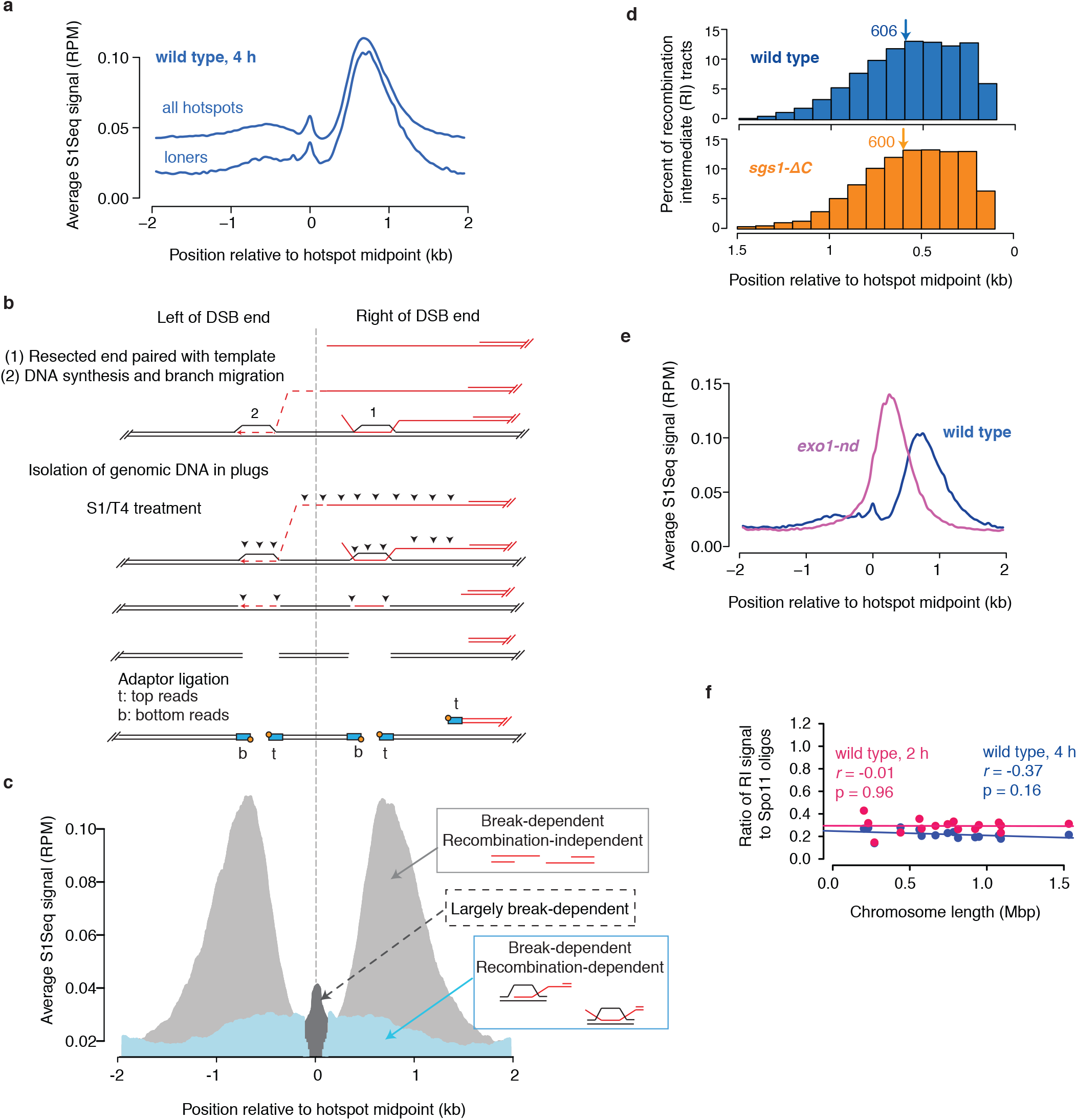
Features of recombination intermediates detected by S1Seq. **a**, S1Seq RI signal is not caused by overlapping resection tracts from nearby hotspots. S1Seq profiles are shown (101 bp smoothing, top and bottom reads co-oriented and averaged) for all hotspots (n=3908) and for hotspots with no other hotspot within 3 kb (loners, n=529). Because of the compact yeast genome and the tendency for many promoters to contain hotspots, resection tracts from neighboring hotspots often overlap, which contributes to presence of S1Seq reads of the “wrong” polarity when considering all hotspots in aggregate. However, overlapping resection tracts are not the only source: wrong-polarity reads are still present even when considering only loner hotspots. **b**, A cartoon to illustrate how RIs might be incorporated into S1Seq sequencing libraries. For simplicity only strand-exchange events involving the right side of a DSB are depicted. We favor the idea that RI signal represents strand-invasion intermediates rather than double-Holliday junctions, because the latter are likely to be resistant to S1 digest. Cleavage by S1 nuclease of ssDNA in D-loops (plus polishing by T4 DNA polymerase) is expected to yield ligatable ends as shown. In scenario 1, a D-loop from invasion of the resected DSB end into a homolog or sister chromatid can yield bottom-strand reads that map to the right of the DSB, i.e., the opposite polarity from a resection endpoint. In scenario 2, D-loop migration could yield top-strand reads that map to the left of the DSB. Note that in each scenario, reads of the “correct” polarity for resection endpoints are also expected to arise from these specific types of RI. Therefore, for quantitative analysis throughout this study we subtracted the bottom-strand reads from the top-strand reads when we examined resection to the right of hotspots, and we subtracted the top-strand reads from the bottom-strand reads when we examined resection to the left. Generally speaking, however, this correction had little effect on final results because of the much higher abundance of the signal for resection endpoints compared with RIs (for more details see Supplementary Methods). **c**, A graphical representation of the different species detected by S1Seq around loner hotspots: resection endpoints (light grey), RIs (blue), and unresected DSBs (dark grey). **d**, Distribution of RI signal relative to hotspot midpoints. Arrows indicate means (bp). The frequency and distribution of RI lengths are identical to wild type in the *sgs1-ΔC* mutant, indicating that the multi-chromatid joint molecules formed at high levels in this mutant^38,39^ are S1 nuclease-resistant. **e**. Genome averages of strand-specific S1Seq signal at 4 h (101-bp smoothing) centered on midpoints of Spo11-oligo hotspots; similar to Fig. 1f except that only loner hotspots are shown. Note that RI signal forms a shoulder to the left of hotspot midpoints in the *exo1-nd* mutant, indicative of shorter spread of the RI branch points. We interpret this behavior as more consistent with events in scenario 1 in panel b, because scenario 2 events might be expected to be independent of resection length. Furthermore, given how much DNA synthesis would be necessary to account for the distribution of events under scenario 2 as well as the lack of observable extension of 3′ DSB ends in vivo^4,33,35^, we favor scenario 1 as the major source of RI signal. **f**. No correlation between chromosome size and the yield of S1Seq RI signal per DSB (Spo11-oligo count). RI signal was measured by summing the regions from –100 bp to –1000 bp (top strand) and +100 bp to +1000 bp (bottom strand) relative to the midpoints of loner hotspots.

**Extended Data Figure 4.**
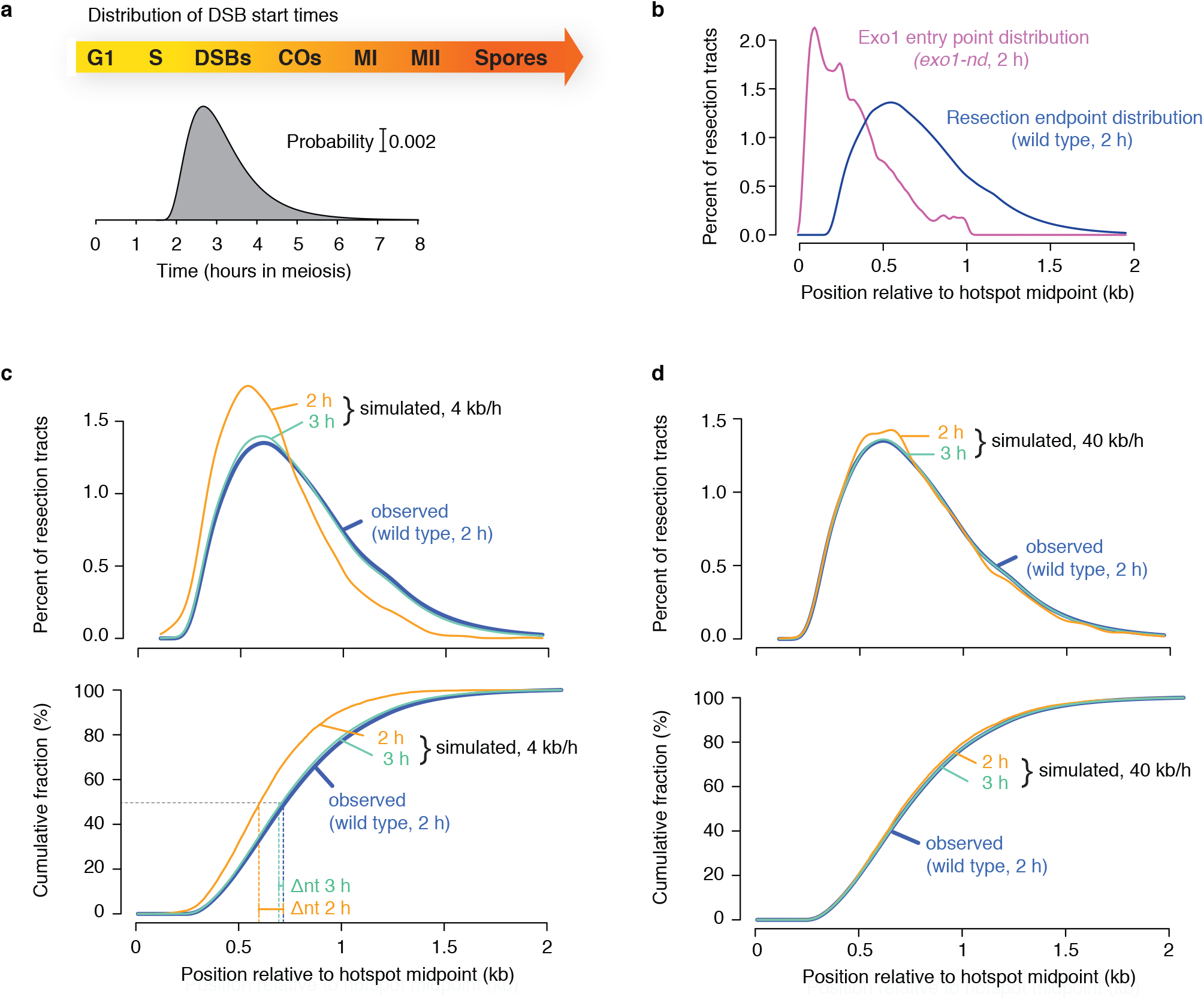
Modeling Exo1 resection speed. **a**, Time distribution of DSB formation by Spo11, based on published DSB entry and exit cumulative curves^33^. **b**, Spatial distribution of Exo1 entry points and endpoints at 2 h in meiosis, namely the *exo1-nd* and wild-type (predicted by shifted geometric model) profiles, respectively (10-bp binned, 11-bin smoothing). **c, d**, Examples of simulations at two different Exo1 rates: 4 kb/h, equivalent to the long-range resection rate in vegetative cells, and 40 kb/h, the rate of Exo1 on naked DNA in vitro. For all the simulations, DSBs were formed in accord with known timing (panel a), were clipped instantaneously by Mre11 to establish Exo1 entry points (panel b), and were allowed to resect *in silico* at 4 kb/h or 40 kb/h to final lengths matching the wild-type distribution (panel b). Simulated endpoints (percent and cumulative fraction) at different times were then compared to the observed endpoint distribution (panels c and d). See also **Supplemental Movies 1 and 2.** ***Notes***: This exercise evaluates the plausibility of specific Exo1 rates but does not directly measure resection speed. At 4 kb/h, simulated tracts were markedly shorter than observed tracts at 2 h, with continuing presence of partially resected DSBs throughout meiosis (panel c)(see also Fig. 4c and Supplemental Movie 1). When the Exo1 rate was 40 kb/h, the value from singlemolecule studies for Exo1 resecting naked DNA^23^, simulated and observed tract lengths matched well throughout the time course (panel d) (see also Fig. 4c and **Supplemental Movie 2**). Another way to conceptualize these results is by calculating the average distance Exo1 needs to resect, which is the difference between the average *exo1-nd* and wild type resection lengths, 822 nt – 373 nt = 449 nt. At 4 kb/h, this would take about 6.75 min whereas at 40 kb/h only about 40 sec. Considering that DSBs are formed asynchronously and continuously for several hours in a sporulating culture, the fact that partially resected DSBs are essentially undetectable at any time point means that near-instantaneous completion of resection is needed to match the observations.

**Extended Data Figure 5.**
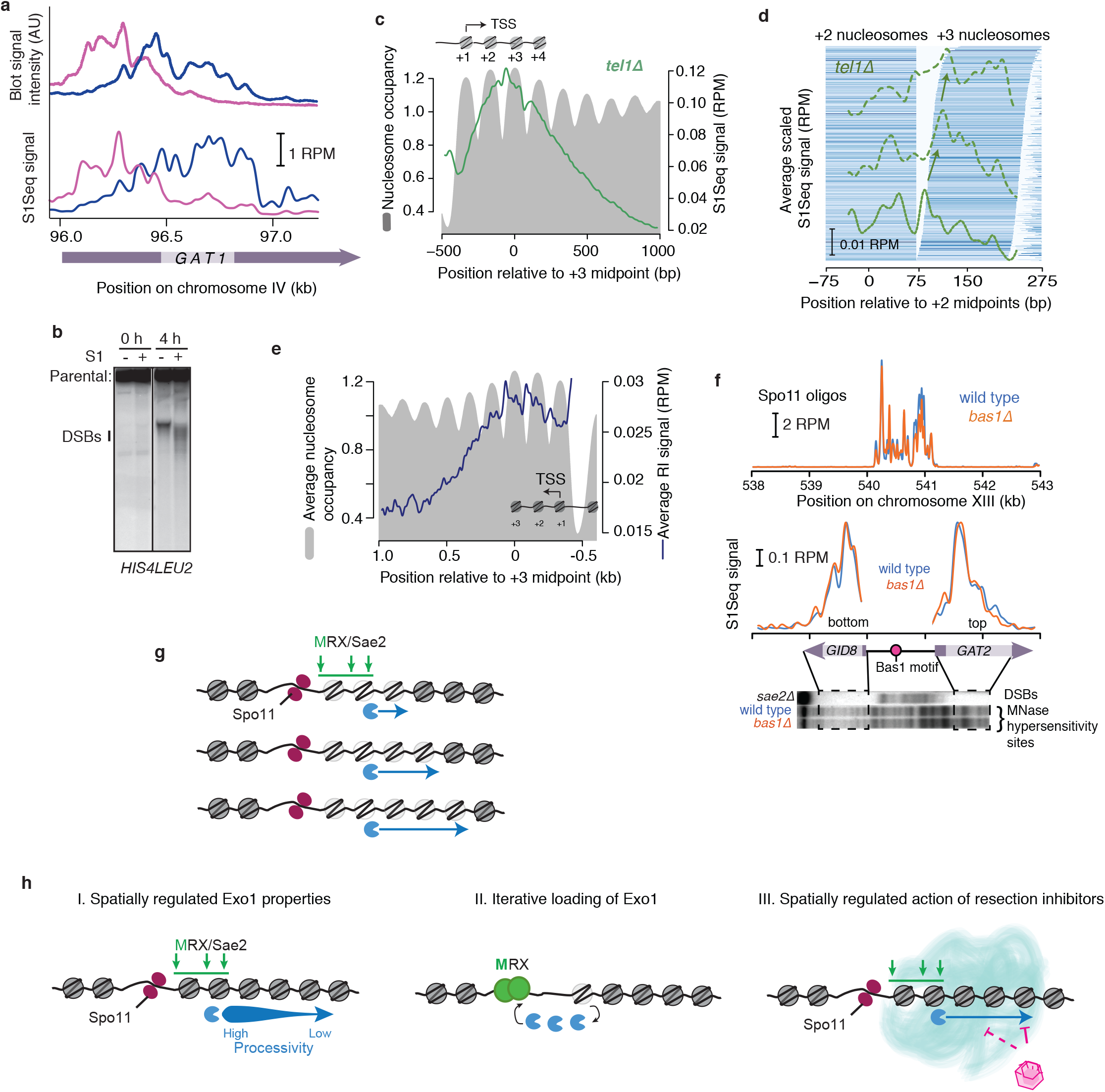
Chromatin structure shapes the resection landscape. **a**, Comparison of Southern blot of S1-treated DSBs with S1Seq maps. The top traces are lane profiles for the 4-h time points from wild type and *exo1-nd* from Fig. 5a; the bottom traces are the S1Seq profiles from the approximately corresponding region. Note that horizontal scales are not precisely matched because of logarithmic mobility of the DNA on the agarose gel. Importantly, the banding pattern did not change if the amount of S1 nuclease was varied (data not shown), so it is not a consequence of incomplete ssDNA digestion. **b**, DSBs at *HIS4LEU2* do not display a discrete banding pattern after S1 digestion. This difference compared with *GAT1* (and, from S1Seq data, most other natural hotspots) presumably reflects an absence of a well-positioned nucleosomal array adjacent to the artificial hotspot. **c,d**, Comparison of *tel1Δ* S1Seq at 4 h with chromatin structure. Data are presented as for Fig. 5c,d around the indicated nucleosome positions (n=1815 for panel c, n=1034 for panel d). **e**, Comparison of RI distribution with chromatin structure. Data are presented as for Fig. 5c except that the wild-type RI S1Seq profile at 4 h is used instead of the resection endpoint distribution. **f**, A counter-example for Fig. 5g,h. Spo11 oligos and a Southern blot of MNase-digested chromatin^26^ are shown in comparison to the S1Seq signal around a hotspot that contains a binding site for Bas1, but for which chromatin structure is unaltered in the *bas1Δ* mutant. S1Seq patterns are likewise unaltered. **g**, Nucleosome remodeling as a principal determinant of resection termination positions. In this schematic, DSBs are shown occuring at the same location in three different cells. Green arrows denote the region where nicking by MRX plus Sae2 occurs, and the blue Pacman and arrow show the extent of Exo1-dependent resection. In this model, formation of each DSB is accompanied by subsequent destabilization or outright eviction of some number of adjacent nucleosomes (translucent gray circles). Given the strong chromatin signature for resection in the absence of Exo1, the hypothetical chromatin remodeling presumably occurs concomitant with or following Mre11-dependent nicking. Exo1 is then able to resect rapidly and processively, but only until it encounters the first intact nucleosome (dark gray circles). In vitro, removing H2A and H2B or replacing H2A with the H2A.Z variant (Htz1) allows modest Exo1 activity^27^, so such partial nucleosome remodeling might be sufficient to facilitate resection in vivo. However, because meiotic resection appears to approach the in vitro rate on naked DNA, we favor instead the possibility that nucleosomes are often fully evicted. Eviction of a minimum of one nucleosome beyond the most distal Mre11-dependent nick might account for the good fit of the shifted geometric model for Exo1 run lengths. Furthermore, the nucleosome remodeling hypothesis reconciles the paradox that resection occurs extremely rapidly even though chromatin appears to be a strong block to Exo1 progression, as judged by the apparent tendency for Exo1 to stop upon nucleosome entry and by the fact that Exo1 run lengths are much shorter in vivo (mean = 446 nt) than in vitro (mean = 5.6 kb; ref. 23). **h**, Additional factors that could account for the apparent shifted geometric distribution of Exo1 run lengths. (I) Changes in the intrinsic properties of Exo1 that govern its ability to digest nucleosomal substrates. For example, a posttranslational modification that increased Exo1 processivity but that decayed stochastically as Exo1 resected away from the DSB could work to ensure a minimal resection length. (II) Iterative loading of Exo1 within a zone close to the DSB could also explain how Exo1 can efficiently resect for a minimum distance. The MRX complex, for example, has been shown to recruit Exo1 *in vivo* and thus may increase its local concentration around the break site^40^–^42^. (III) Constrained access of inhibitors could influence the distribution of termination sites. The pink polygon represents an Exo1 inhibitor that is itself inhibited within a zone close to the DSB (indicated by the teal cloud). Known inhibitors of Exo1 include Rad9, Rad53 and rpa^23,41,43–46^. In all of these models, activation of Tel1 at the DSB could provide a plausible means for spatially patterning the relevant modifier of Exo1 activity.

## Methods

### Yeast Strains and culture methods

All strains used in this study were SK1 background^47^ and the genotypes are listed in Extended Data Table 2. The *sgs1-ΔC795* allele was provided by N. Hunter and the *exo1-D173A* allele by R. Liskay^15^. The *dmcl and tell* deletions were made by replacing the coding sequence with the hygromycin B phosphotransferase gene *(hphMX4)*. Gene disruption was verified by Southern blot. The *bas1* and *ino4* deletion alleles were made as described^26^. Diploid cells were streaked from a frozen stock onto YPD plates and allowed to grow at 30°C for two days. A single colony was inoculated into 20 ml liquid YPD medium and grown for >24 hr at 30°C. The saturated YPD culture was used to inoculate the appropriate volume of YPA medium (1% yeast extract, 2% Bacto peptone, 2% potassium acetate, 0.001% antifoam 204 (Sigma)) to OD_600_ 0.2 and grown in 2.8 l baffled Fernbach flasks at 250 rpm at 30°C for 14 hr. Cells at indicated time points (75 ml for 0 and 2 h, 66 ml for 4 h, 60 ml for 6 h and 54 ml for 8 h) were harvested, washed with 50 mM EDTA pH 8.0 and stored at -80°C until plug preparation.

### Preparation of DNA and sequencing libraries for S1Seq

Genomic DNA was extracted in low-melting point (LMP) agarose to yield intact high molecular weight DNA. Each cell pellet was resuspended in 150 μl 50 mM EDTA pH 8.0 and allowed to thaw at 40°C. 498 μl LMP agarose (2% in 50 mM EDTA pH 8.0) was mixed with 102 μl Solution 1 (SCE (1 M sorbitol, 0.1 M sodium citrate, 60 mM EDTA pH 7.0) plus 5% β-mercaptoethanol and 1 mg/ml zymolyase 100T) and the mix was prewarmed at 40°C before adding to the cells. The resuspended cells were pipetted into plug molds and chilled at 4°C for 30 min. Once solidified, the agarose plugs (three at a time) were expressed into 3 ml of Solution 2 (0.45 M EDTA pH 8.0, 0.01 M Tris-HCl pH 7.5, 7.5 % β-mercaptoethanol, 10 μg/ml RNase A) and incubated at 37°C for 1 h. Solution 2 was then replaced with Solution 3 (0.25 M EDTA pH 8.0, 0.01 M Tris-HCl pH 7.5, 1% SDS, 1 mg/ml proteinase K) and plugs were incubated at 50°C overnight. Plugs were finally washed three times with 3 ml 50 mM EDTA pH 8 and stored in plug storage solution (0.05 M EDTA pH 8.0, 50% glycerol) at -20°C until all plugs were ready to be processed for library preparation.

For efficient in-plug overhang removal and adaptor ligation, a sequential treatment with S1 nuclease and T4 DNA polymerase was required before ligation. Even though S1 nuclease treatment was sufficient to remove the overhang tails, the subsequent ligation step was not efficient, unless a clean-up reaction with T4 DNA polymerase was performed (E.P.M., unpublished observations). Ten plugs of each sample to be analyzed were equilibrated in 500 μl 1 × S1 buffer (50 mM sodium acetate pH 4.5, 0.28 M NaCl, 4.5 mM ZnSO_4_) per plug in 2 ml conical tubes for 30 min and this was repeated three times. Fresh 1 × S1 buffer (500 ul) containing 9 U of S1 nuclease (Promega) was added to each tube and the plugs were incubated on ice for 15 min to allow the enzyme diffuse into the plugs, followed by 20 min incubation at 37°C. S1 nuclease was inactivated by addition of EDTA pH 8.0 to a final concentration 10 mM and incubation on ice for 15 min. Plugs were rinsed with 1 × TE and further equilibrated in 500 μl T4 polymerase buffer (1 × T4 DNA ligase buffer (NEB), 1 × BSA, 100 μM dNTPs), four times for 30 min each. Fresh T4 polymerase buffer (500 μl) containing 30 U of T4 DNA polymerase (NEB) was added to each tube and the plugs were incubated on ice for 15 min, followed by 30 min incubation at 12°C. T4 polymerase was inactivated by addition of EDTA pH 8.0 to a final concentration of 10 mM and incubation on ice for 15 min. At this point all buffer was aspirated, the plugs were rinsed with 1 × TE and incubated at 75°C for 20 min to fully inactivate T4 polymerase. Plugs were allowed to gradually cool and solidify, before equilibrating in 250 μl 1× T4 ligase buffer (NEB), four times for 15 min each. After the last equilibration step, 200 μl of the buffer were removed leaving each plug immersed in 50 μl 1 × T4 DNA ligase buffer, to which 1 μl of 50 μM adaptor and 1 μl of 2000 U/μl T4 DNA ligase were added. Ligation took place at 4°C for ≥18 h. The adaptor was prepared the day of the experiment by mixing oligos P5-top (5BiosG/ACACTCTTTCCCTACACGACGCTCTTCCGATCT, biotin attached to the 5′-end of the oligo) and P5-bottom (5Phos/AGATCGGAAGAGCGTCGTGTAGGGAAAGAGTGT/3InvdT, 5′-end phosphorylated with inverted dT incorporated at the 3′-end to block ligation) at equimolar concentration, boiling for 5 min and cooling down at room temperature for at least 1 h.

To retrieve the DNA from the agarose plugs, the Epicentre GELase Enzyme Digestion protocol was used, followed by phenol extraction and ethanol precipitation. During this process plugs of DNA from a single culture and time point were pooled. The purified DNA was then subjected to size selection by electrophoresis through a 1% LMP 1× TAE agarose gel at 80 V/cm for 2 h to separate the high-molecular-weight genomic DNA from the excess unligated adaptor. The genomic DNA was excised and extracted from the gel using the Epicentre GELase enzyme protocol, phenol extraction and ethanol precipitation; then the DNA was dissolved in 100 μl 1 × TE. To ensure complete removal of unligated adaptor, each sample was further purified by passing three times through Chroma Spin-1000 (Clontech) columns. The eluates were then subjected to shearing according to the Covaris instrument protocol to DNA fragment sizes ranging between 200-500 bp.

Following shearing, fragments containing the biotinylated adaptor were enriched by affinity purification with streptavidin. For each sample, 50 μl of streptavidin M-280 beads (Roche) were used, prewashed twice with 1 × TE and twice with 1 × B&W (binding and washing buffer, 10 mM Tris-HCl pH 7.5, 1 mM EDTA, 2 M NaCl), according to the manufacturer’s instructions. Binding was carried out at 20°C for 30 min, followed by two washes with 500μl 1× B&W buffer and two washes with 500 μl 10 mM Tris-HCl pH 7.5. The remaining steps were performed with the DNA fragments still bound to the streptavidin beads. Because sheared ends are not always readily ligatable, an end-repair step was included according to Epicentre’s End-it DNA Repair kit protocol. The beads for each sample were incubated in 100 μl of reaction mix according to manufacturer’s instructions and incubated at 20°C for 45 min. Following washes with 500 μl 1× B&W buffer (twice) and 500 μl 10 mM Tris-HCl pH 7.5 (twice), the beads were incubated in 100 μl of 1 × T4 DNA ligase buffer containing 400 U T4 ligase and 1 μM P7 adaptor. The adaptor was prepared by annealing oligos P7 top (5Phos/GATCGGAAGAGCACACGTCTGAACTCCAGTCAC/3InvdT) and P7 bottom (GTGACTGGAGTTCAGACGTGTGCTCTTCCGATC) at equimolar concentration, boiling for 5 min and allowing to cool down at room temperature for at least one hour. Ligation of the P7 adaptor was allowed at 20°C for 4 h and was followed by washes with 500 μl 1 × TE and 500 μl 10 mM Tris-HCl pH 7.5 (three times each). Beads of each sample were resuspended in 20 μl 10 mM Tris-HCl pH 7.5.

Each sample of beads was split into two PCR reactions in a total of 50 μl containing 10 μl of the resuspended beads, 1 × Phusion HF buffer, 0.2 mM dNTP, 1 U Phusion HF polymerase (Thermo Scientific), 0.4 μM P5 universal primer (AATGATACGGCGACCACCGAGATCTACACTCTTTCCCTACACGACGCTCTTCCGATCT) and 0.4 μM P7 indexed primer (CAAGCAGAAGACGGCATACGAGATCGTGATGTGACTGGAGTTCAGACGTGTG, shown here with Illumina TruSeq index 1 underlined). PCR was initiated by a denaturation step at 98°C for 30 sec, followed by 16 cycles of amplification (98°C for 10 sec, 65°C for 20 sec, and 72°C for 20 sec). PCR products were pooled, and the DNA was precipitated by adding ammonium acetate to 2.5 M and 2.5 volumes of ice-cold 100% ethanol. Following an overnight incubation at −20°C, the samples were spun at 16000 g for 30 min, washed with ice-cold 70% ethanol, air-dried and dissolved in 20 μl 10 mM Tris, pH 7.5. PCRs from each sample were pooled and separated on a 5% non-denaturing polyacrylamide gel in 1 × TBE. The gel piece between 200–600 bp was excised, crushed, and eluted in 300 μl 10 mM Tris pH 8.0 at 37°C overnight. The elution mixture was spun through a SPIN-X column, and DNA was precipitated with ammonium acetate and ethanol as above. The DNA pellet was dissolved in 20 μl 10 mM Tris pH 7.5 and sequenced on the Illumina HiSeq platform (50 bp single-end) in the Integrated Genomics Operation at Memorial Sloan Kettering Cancer Center.

While we were optimizing this protocol, a similar method designed to map DSBs at high spatial resolution, BLESS, was described^48^. Another method, called EndSeq, has also been recently described^49^. A key difference between S1Seq and both BLESS and EndSeq is our use of S1 nuclease, which we found necessary to efficiently remove long stretches of ssDNA. In addition, BLESS uses a formaldehyde fixation step which we do not use, and which we speculate might cause unwanted damage to the DNA samples.

### Physical analysis of resected and blunt-ended DSBs

For direct detection of DSB fragment migration before and after ssDNA removal at *GAT1*, the genomic DNA isolated in plugs was subjected to restriction endonuclease digestion, gel electrophoresis and Southern blot analysis. The plugs were treated with S1 endonuclease as described above but instead of T4 DNA polymerase and T4 DNA ligase treatment they were digested with PstI-HF (NEB) in plugs equilibrated in the appropriate restriction enzyme buffer as described^13^. Digested DNA was separated on 0.8% agarose gels in 0.5× TBE with electrophoresis at 80 V/cm for 16 h and re-circularization of the buffer, then detected by Southern blot hybridization using a ^32^P-labeled DNA fragment adjacent to one of the PstI sites as a probe. The primer sequences for amplification of the probe were 5′-CGCGCTTCACATAATGCTTCTGG and 5′-TTCAGATTCAACCAATCCAGGCTC.

### Bioinformatics analysis

Mapping of the reads onto the S288c reference genome (SacCer2) was performed using the SHRiMP mapper^50^ (gmapper-ls) with arguments:

> -E -U -n 1 -Q --sam-unaligned --strata -o 10001 -N 20

Before mapping, adaptor sequences were removed using fastx_clipper (http://hannonlab.cshl.edu/fastx_toolkit/) and a custom script. Code used for read processing and mapping is available online at https://github.com/soccin/S1Seq. After mapping, the reads were separated into unique and multiple-mapping sets, but only uniquely mapping reads were analyzed in this study.

Analyses were performed using the R package (RStudio version 0.99.879, R version 3.1.2). Before any analysis, maps were curated by masking (set to NA) 500 bp at the ends of each chromosome, because telomeres represent naturally occurring resected DNA ends and therefore give a high frequency of reads. We further masked regions that gave a high frequency of meiotic DSB-independent reads, defined as positions with more than 10 reads in the high-depth biological replicates and/or more than 5 reads in the lower depth biological replicates from the 0-h maps from wild type, *spo11-Y135F, dmc1Δ*, and *exo1-nd* plus the 4-h map from *spo11-Y135F*. A table of mask coordinates is provided at https://github.com/soccin/S1Seq. Moreover, reads mapping to mitochondrial DNA or the 2 μ plasmid were excluded. Each map was then normalized to reads per million remaining mapped reads, then biological replicates were averaged. The Gene Expression Omnibus (GEO) (http://www.ncbi.nlm.nih.gov/geo) accession number for the raw and processed sequence reads is pending.

All analyses were performed using a recent hotspot list compiled from a combination of multiple independent wild-type Spo11-oligo maps^51^. For the S1Seq resection and recombination lifespan analyses, the left arm of chromosome III was censored to avoid inconsistencies because some strains had the *HIS4LEU2* and *leu2:hisG* artificial hotspots on this chromosome arm. Published maps of nucleosome occupancy^13^ and histone H3 midpoints^52^ were used. We defined subtelomeric hotspots (n=60) as those residing in the 20-kb telomere-proximal region; pericentromeric hotspots (n=82) are those residing in the 20-kb zone surrounding centromeres. For all modeling analyses, resection length histograms, resection length calculations, resection endpoint distributions centered on nucleosome midpoints, and analyses of RIs, we used the subset of hotspots for which no other hotspot was located within 3 kb, and for which hotspot width was less than 400 bp (n=405). S1Seq reads of polarity opposite to expectation for resection endpoints were subtracted to correct for RI signal. Adjustment for RIs had only a small effect on calculated values. For example, wild-type mean resection length was 779 nt without RI subtraction and 822 nt with subtraction, and the shift required for best fit in the shifted-geometric modeling experiment was 260 nt without RI subtraction versus 220 nt with subtraction.

### Modeling Exo1 resection run lengths in wild type

We built a model based on the assumption that Exo1 starts resecting at Mre11-dependent nicks or gaps, whose distribution is measured by the resection maps from the *exo1-nd* mutant. We also assumed that a single Exo1 molecule resects DNA from 5′ to 3′ until it dissociates from its substrate without chance to rebind. If there is an equal probability of dissociation at each nucleotide step, Exo1 resection tract lengths will follow a geometric distribution determined by that stepwise dissociation probability. Note that application of this model to S1Seq data yields an estimate of the average apparent probability of resection termination at each step. We do not exclude the possibility of there being considerable microscopic variation in this probability at different locations in the genome or between individual resection events in the population.

After subtraction of RI signal from resection endpoint signal (see above), resection profiles for loner hotspots were averaged and binned (10 bp) to make the analysis computationally tractable. Moreover, the signal close to hotspot midpoints was excluded from the analysis by setting values of positions <200 bp for wild type and <20 bp for *exo1-nd* to zero. Finally, an estimated background was removed by subtracting from all values the value of signal 2 kb away from hotspot midpoints.

Let *M*be the observed *exo1-nd* mutant resection endpoint distribution in 10-bp bins: *M* = {*m*_0_, *m*_1_,…, *m*_200_}. Similarly, let *W* be the binned wild-type resection endpoint distribution: *W* = {*w*_0_, *w*_1_,…, *w*_200_}. Thus, *m_i_* and *w_i_* are empirical estimates of how many resection endpoints are located between (10*i* – 5) bp to (10*i* + 4) bp away from hotspot midpoints. We only considered signals located within 2 kb of hotspot midpoints (i.e., max *i* = 200). Let *R* be the distribution model for the Exo1 resection run lengths, *R* = {*r*_0_, *r_l_*,…, *r*_200_}, where *r_k_* is the probability that Exo1 stops resecting 10k nucleotides away from the Exo1 entry sites. We empirically chose 2 kb (i.e., max *k* = 200) as the resection length limit based on the average wild-type resection signal around hotspots.

Let *θ* be the unknown parameter of a geometric distribution for *R* and let *L*(*θ*) = {*l*_0_, *l*_2_,…, *l*_200_} be the likelihood function of *θ*. We wished to estimate *θ* from the combination of the *exo1-nd* mutant and wild-type data. The wild-type resection endpoint frequency at a given distance from hotspot midpoints is the sum of the resection tracts that start anywhere closer to hotspot midpoints and that end at that given distance. Thus *L* can be described as a series of linear equations combining *M*, *W*, and *R*:

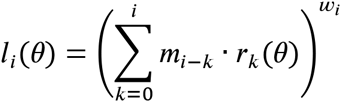

We optimized *θ* to maximize the log likelihood using the optimize function of R (*θ* = 0.00200). As an alternative, we considered a shifted geometric distribution *R*′ for Exo1 run lengths, which can be described as:

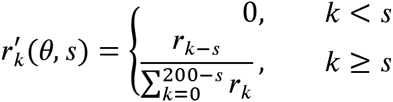

where *s* is the number of 10-nucleotide bins by which to shift the geometric distribution. We tested shifts of 0–30 bins (i.e., 0–300 nucleotides) and optimized *θ* and *s* to maximize the log likelihood (*θ* = 0.00440, *s* = 22). The Exo1 run length distributions predicted by the geometric and shifted geometric models with these best-fit parameters are presented in the middle panel of Fig. 4a.

### Simulating Exo1 resection speed

To evaluate possible rates for resection by Exo1, we performed Monte Carlo simulations to generate populations of resected DNA molecules as a function of time in meiosis, and compared them to observed resection distributions. Details are provided below, but an overview is as follows. In the simulations, a total of 1 million DSBs were allowed to form over the course of meiosis with timing similar to published time course data^33^. This procedure accommodates the fact that DSBs form in a semi-synchronous manner over several hours in meiosis. These DSBs were then assumed to be clipped instantaneously by MRX plus Sae2, with positions sampled from the empirical distribution of resection endpoints in the *exo1-nd* mutant. The assumption of instantaneous clipping biases the model in favor of slower Exo1 speeds, i.e., by having the clipping step take no time, Exo1 thereby is allowed more time to resect to completion. We further sampled to establish Exo1 run lengths using the best-fit shifted-geometric model. Given these three sampled values for each DSB (time of formation, position of Exo1 entry point, and distance traveled by Exo1), the spatial distribution of resection endpoints can be calculated arithmetically at any given time in meiosis as a function of resection speed. By comparing the simulated resection profiles to observed resection profiles, we can evaluate the plausibility of a given Exo1 speed, i.e., whether that speed is fast enough to recapitulate empirical resection lengths.

To establish a model for DSB timing in meiosis, we started with published analysis of DSB formation at the *HIS4LEU2* hotspot (Time Course 36)^33^, in which DSBs formed from 1.5 h to 8 h after transfer to sporulation medium. From these data, a normal distribution fit well to the time profile of DSB formation, but we considered that a model for global DSB timing would be likely to display a positively skewed DSB time distribution given temporal differences between different parts of the genome^53^ and given cell-to-cell variability in meiotic prophase length. Therefore, we used a shifted log normal distribution as a DSB time distribution (Extended Data Fig. 4a). However, we reached the same overall conclusions about plausibility of resection speeds if we used a normal distribution for the DSB timing profile instead (data not shown).

To define the distribution of Exo1 entry points, we used the *exo1-nd* resection endpoints from 2 h, with the rationale that a significant number of DSBs have already formed at this point, but there has not been sufficient time for DSBs to disappear via recombination. To define Exo1 endpoints, we used the shifted geometric run-length model fitted as described above, except using the data at 2 h in meiosis *(θ* = 0.00382, s = 16). Note that, because total resection length in wild type is slightly shorter at 2 h than at 4 h (Fig. 2d), choosing to use input data from the 2 h time points also biases our simulations in favor of slower Exo1 speeds.

To carry out the simulations, we generated a random sample of 1 million DSB formation times (in min after meiosis induction) using the rlnorm function in R with log mean = 4.5, log standard deviation = 0.5 and shift = 1.5 h, taking into account values of time points ≥1.5 h and ≤8 h. A random sample of 1 million Exo1 start points was generated by sampling with replacement from 10-bp bins between 0 and 2 kb from hotspot midpoints with sampling weights matching the empirical distribution of resection endpoints from the 2-h *exo1-nd* map (Extended Data Fig. 4b). A random sample of 1 million Exo1 run lengths (also in 10-bp bins) was generated from the shifted geometric model with parameters listed above.

From these randomly generated values, the distribution of resection endpoints can be calculated directly at any time in meiosis (*t*) for any given Exo1 speed (*v*). If *τ_i_*, *μ_i_* and *ε_i_* are the sampled values for formation time, Mre11-dependent clipping position, and Exo1 run length, respectively, for the *i*th simulated DSB, then the resection endpoint distance *d_i_* as a function of time in meiosis can be described as:

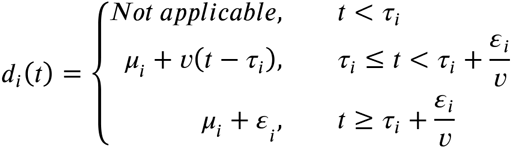

Results of these simulations are shown in **Supplemental Movies 1 and 2** for resection speeds of 4 kb/h and 40 kb/h, respectively. A copy of the R code for carrying out the simulations is provided at https://github.com/soccin/S1Seq.

---

**Extended Data Table 1.**
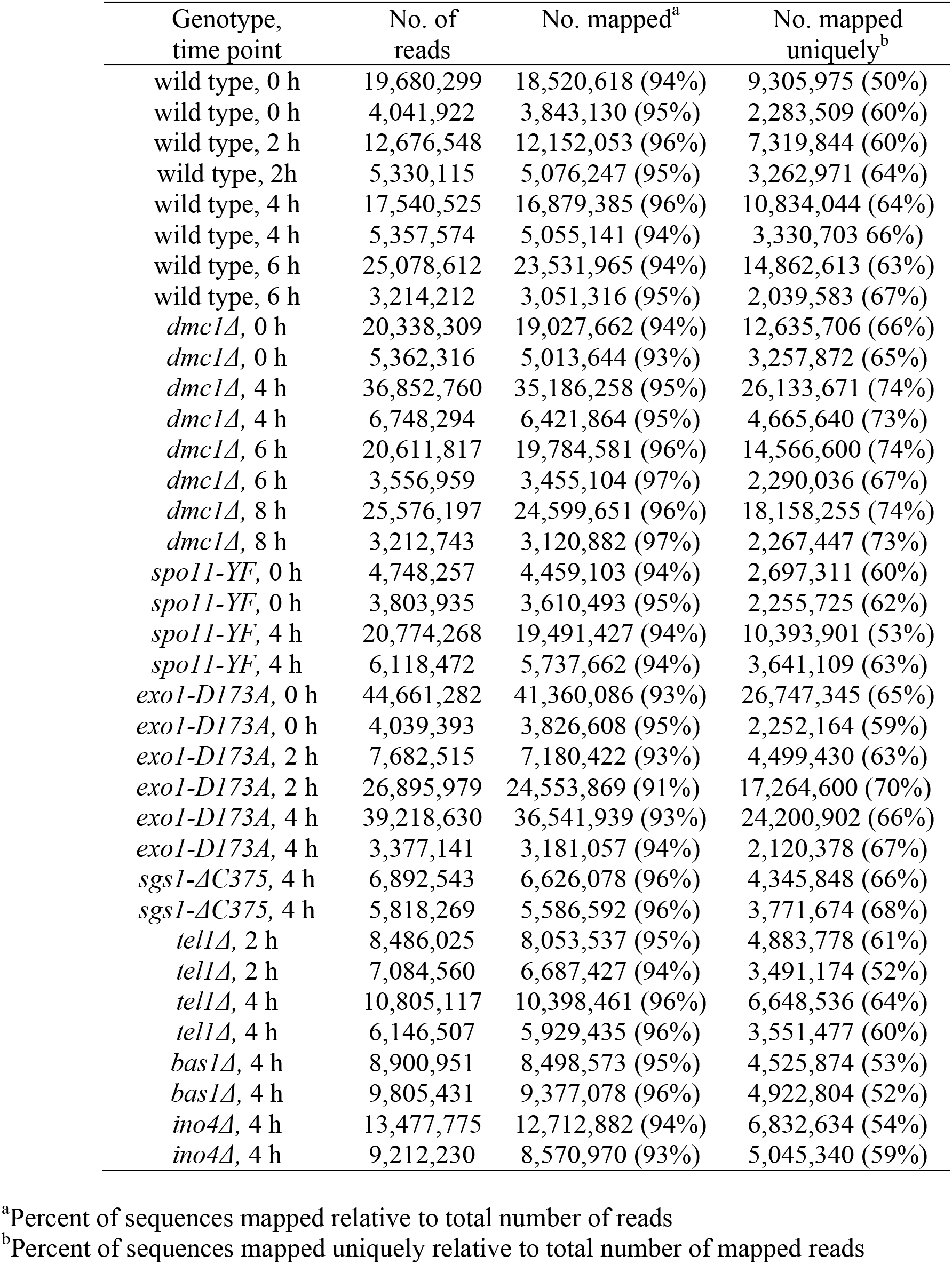
S1Seq mapping statistics

**Extended Data Table 2.**
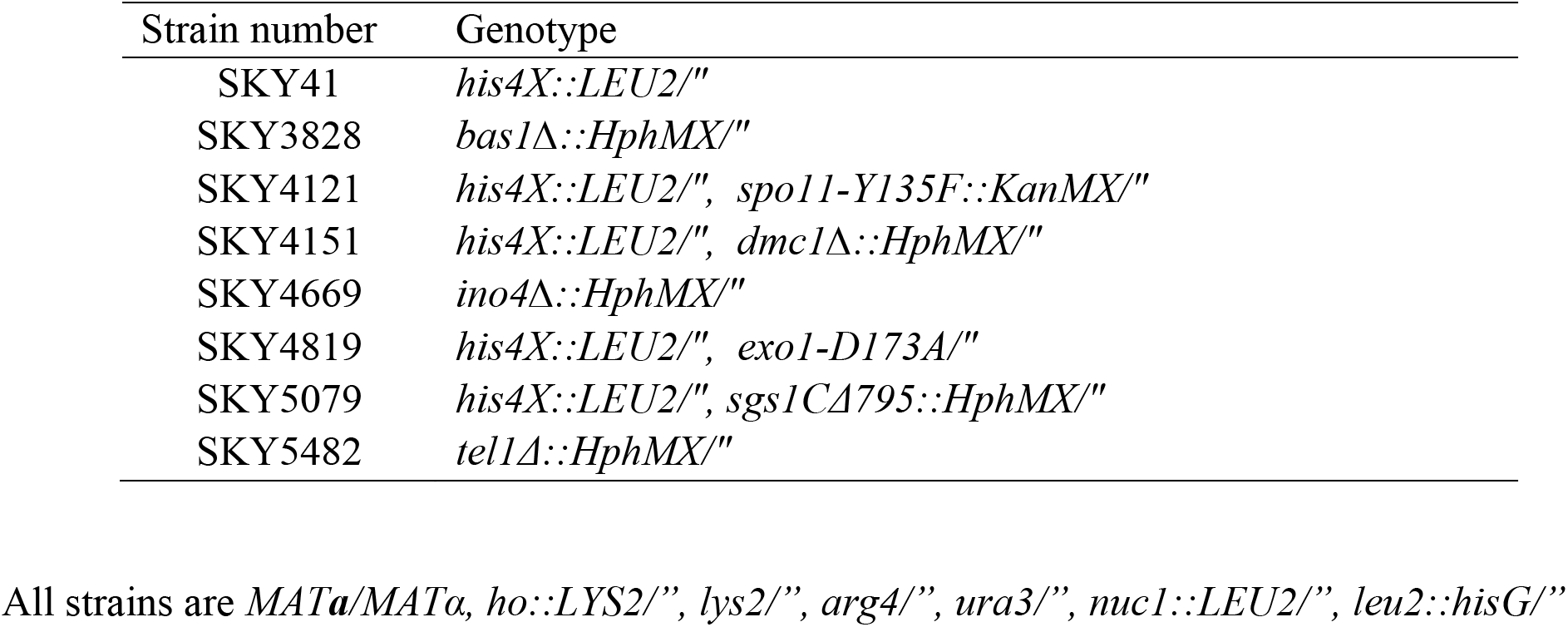
Yeast Strains.

## Supplementary Discussion

A number of studies in the past decade have approached the question of whether and how chromatin structure is altered in the vicinity of a DSB, and how such changes influence DSB processing and repair^29,31,54,55^. However, controversy surrounds the subject of histone eviction before, during or after DSB end processing.

Several chromatin immunoprecipitation (ChIP) studies in yeast have monitored histone loss around the site of an unrepairable DSB. One of the earliest such studies reported that there was no detectable loss of histone H2B in a 5-kb window around the DSB by 60 min after break induction, when approximately 20% of the DSB-containing chromosomes had been resected for at least 0.9 kb^56^. In contrast, another study monitoring H2B and H3 ChIP in a 6-kb window around the inducible DSB reported loss of both histones that was particularly apparent 90 min after break induction^30^. A caveat in the latter study was the lack of normalization of the ChIP signal to the DNA input signal, thus a decrease of histone ChIP signal due to histone loss could not be distinguished from loss of DNA due to end resection. Intriguingly, however, even when later studies applied such a normalization step, conflicting results were reported. Gasser and colleagues observed an approximately two-fold decrease in H3 ChIP signal within 1 kb of the DSB; this decrease was largely dependent on the INO80 chromatin remodeling complex and was most apparent 2–4 h post DSB induction, with no histone loss detected at sites >5 kb away^28^. On the other hand, the Tyler group found that loss of H3 ChIP signal 0.6 kb away from the DSB closely mirrored the loss of DNA input signal over time, suggesting that the rate of chromatin disassembly around the DSB is tightly coupled to the rate of end resection and that there is no histone loss prior to resection^57^. A more recent study by the Ira group reached a similar conclusion^43^.

One potential basis for the differing results is the use of ChIP as an experimental read-out^29^. The efficiency of formaldehyde cross-linking of histones to DNA may vary between laboratories. Moreover, because crosslinking efficiency differs significantly depending on whether the DNA is double-stranded (before resection) or single-stranded (after resection), there are complications in interpreting ChIP data solely in terms of retention or loss of histones.^58^ Also, it has been pointed out that in studies where histone loss around a DSB site has been detected, the reduction was usually two-to four-fold, whereas the transcriptional histone eviction leads to much greater (five- to ten-fold) decreases in ChIP signal^29^.

Additional lines of evidence for chromatin structure changes around DSBs come from analysis of ATP-dependent chromatin remodelers. Genetic studies in *S. cerevisiae* have demonstrated an important role for the RSC and INO80 ATP-dependent chromatin remodeling complexes in recruitment of MRX to DSBs and in early resection steps, and have implicated the SWI/SNF-like chromatin remodeler Fun30 in extensive (>5 kb) resection^28,43,59–61^. Similar roles have been observed for the mammalian INO80 complex and SMARCAD1, the human Fun30 ortholog^59,62^, although it should be noted that available data in mammalian cells rely on indirect read-outs for resection, typically formation of cytologically observable deposits of the ssDNA binding protein RPA on chromatin.

Finally, studies using MNase hypersensitivity profiles around an inducible DSB in yeast have reported rapid nucleosome migration to form histone-free DNA of a few hundred base pairs immediately adjacent to the break^61,63^.

Taking all of this into account, it appears that chromatin is significantly remodeled around a DSB, but the extent to which histone eviction per se occurs is still a subject of debate and new approaches are necessary to address this question. Furthermore, much of the available data is unable to distinguish between observed chromatin remodeling occurring before (and being required for) DSB resection as opposed to remodeling being a consequence of resection. Moreover, no published studies to our knowledge address these questions in the context of meiotic DSB resection. Our study, using a fundamentally different methodology, provides strong support for a model in which nucleosome destabilization or eviction occurs prior to DSB processing by Exo1 and thereby shapes the resection landscape.

